# Repeated regeneration prolongs the juvenile stage and extends lifespan in the annelid *Platynereis dumerilii*

**DOI:** 10.64898/2026.11.16.767108

**Authors:** Rannyele P. Ribeiro, Kaitlyn Hong, Alina Krawczynski, Faith Stemmler, B. Duygu Özpolat

**Affiliations:** Washington University in St. Louis, St. Louis, United States of America

## Abstract

Regeneration restores tissues lost to injury, but its consequences may extend beyond local repair. Whether regeneration produces lasting systemic effects on developmental timing and lifespan remains poorly understood. Here we investigated the long-term consequences of repeated regeneration in the marine annelid *Platynereis dumerilii*. To determine these consequences, we repeatedly amputated juveniles at monthly intervals, removing approximately half of the body after each regenerative cycle, and compared them with control unamputated siblings. Repeatedly regenerating worms lived approximately 1.5-fold longer than controls and showed delayed sexual maturation, including delayed gametogenic progression at the morphological and transcriptional levels. Somatic progression was also delayed, as indicated by slower accumulation of pigmentation associated with developmental progression, while regenerative competence was maintained across amputations. To investigate whether regeneration engages distant tissues, we compared head transcriptomes from amputated and unamputated worms. Posterior amputation altered head gene expression within 2 hours, including increased expression of genes associated with younger head profiles, such as *igf-like* and *alpha-tubulin*. Together, our results show that repeated regeneration not only restores lost tissues but prolongs the juvenile state, delays reproductive and somatic maturation, and extends lifespan, a phenomenon we term the Highlander effect. These findings suggest that regeneration may influence organismal life history beyond tissue replacement, with neurohormonal signaling as a potential mediator.

## Introduction

Regeneration has been defined as a successful outcome, where an organism restores lost cells, tissues, or body parts. Yet injury can trigger responses far beyond the tissue that regenerates. For instance, in planarians, wound signals propagate across the body and activate responses in tissues distant from the injury (Fan et al., 2023). Similarly, limb amputation in axolotls activates cell proliferation in uninjured tissues, including the contralateral limb and internal organs (Johnson et al., 2018; Payzin-Dogru et al., 2025). In *Drosophila*, damaged imaginal discs release systemic signals that delay metamorphosis and slow the growth of undamaged organs (Colombani et al., 2012; Garelli et al., 2015). These observations suggest that regeneration is not solely a local morphogenetic process, but an organism-wide physiological event. Whether these systemic effects end when the missing structure is restored or persist to alter the subsequent life trajectory of the animal remains largely unknown (Poss & Tanaka, 2024).

Understanding these systemic effects of regeneration is particularly relevant when injury-regeneration occurs repeatedly. Recurrent tissue loss and replacement could impose cumulative physiological costs, but could also repeatedly engage programs involved in growth, repair and somatic maintenance that may have beneficial outcomes. For example, repeated injury extends lifespan in the asexual annelid *Paranais litoralis* (Martínez, 1996), and regeneration in aging sexual planarians can reverse age-associated cellular, molecular and physiological changes in both regenerated and uninjured tissues (Dai et al., 2025). Whether repeated regeneration can extend life span, and the mechanisms through which this might occur, remains unknown.

The annelid *Platynereis dumerilii* provides an opportunity to address this gap. In *P. dumerilii,* regeneration, growth and reproduction are closely coupled throughout the life cycle. *P. dumerilii* progresses from embryo to larva and juvenile, followed by a metamorphic transition associated with sexual maturation, a rapid post-metamorphic decline and death (Fig. 1A). The juvenile phase is typically the longest and is characterized by continuous growth through the addition of segments from a stem-cell-rich posterior growth zone (PGZ) (Fischer et al., 2010; Gazave et al., 2013). Regenerative capacity is largely associated with this juvenile period and progressively declines as animals approach sexual maturity (Álvarez-Campos et al., 2023; Metzger & Özpolat, 2024; Planques et al., 2019). Growth, regeneration and reproductive maturation are further interconnected through neuroendocrine regulation: brain hormonal activity promotes growth and posterior regeneration while suppressing sexual maturation (Hauenschild, 1955, 1960, 1966; Schenk et al., 2016). Regeneration therefore takes place within the same complex regulatory networks that govern progression toward reproduction.

**Figure 1.**
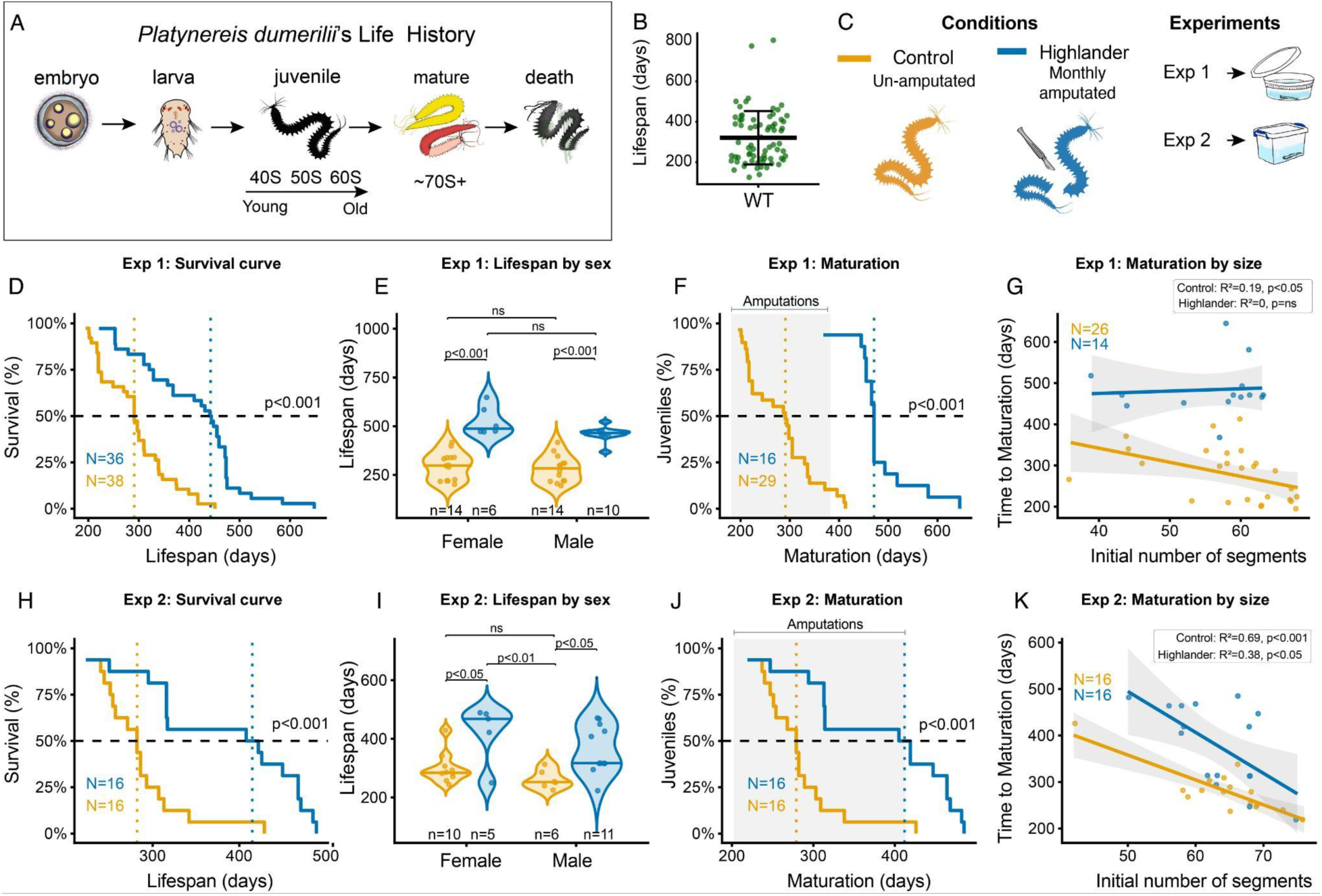
Repeated regeneration extends lifespan and delays sexual maturation. (A) Life history of *P. dumerilii*. (B) Average lifespan of wild-type *P. dumerilii* in laboratory cultures from 2021 to 2023 was 321.1 ± 132.2 days (see Methods for details). (C) Experimental setup. (D–G) Results from Experiment 1, using a sibling cohort born on August 18, 2023, and maintained in round culture boxes: lifespan curve (D), maturation curve (E), relationship between maturation and initial segment number (F), and lifespan comparison by sex (G). (H–K) Corresponding analyses for Experiment 2, using a sibling cohort born on January 29, 2025, and maintained in square culture boxes. Light gray shading in panels E and I indicates the experiment-specific amputation windows. Additional data are presented in Supplementary Fig. S1A–G.

Here we show that repeated posterior regeneration delays developmental progression and extends lifespan in *P. dumerilii*. Using longitudinal measurements of lifespan and development, together with analyses of regenerative competence, germline progression and gene expression, we find that the consequences of repeated regeneration extend far beyond the tissue being replaced. In particular, posterior injury rapidly induces neuroendocrine changes in the head, a region essential for regenerative competence in this species. We therefore refer to this phenotype as the Highlander effect, inspired by the fictional immortals of *Highlander*, who can die only when beheaded. Our results reveal that regeneration can trigger responses far from the wound and produce long-term organism-wide beneficial effects.

## Results

### Repeated regeneration extends lifespan

In *P. dumerilii*, sexual maturation marks a critical life-history transition: upon maturation regenerative competence is lost, reproduction begins, and death follows (semelparity) (Fig. 1A). To determine whether repeated regeneration alters life-history progression in *P. dumerilii*, we examined the effects of repeated amputation-regeneration cycles on sexual maturation and lifespan. We conducted two independent experiments comparing sibling worms randomly assigned to Control or Highlander conditions. In each experiment, Highlander worms underwent monthly amputations removing the posterior half of the body, continuing until all surviving Control worms matured (Fig. 1C; see Methods).

Consistent with the median lifespan of 286 days observed in our cultures (Fig. 1B), Control worms in Experiment 1 had a median lifespan of 291 days (n = 38) (Fig. 1D). In striking contrast, Highlander worms (n = 36) exhibited a significant 1.52-fold extension in lifespan, reaching a median of 442.5 days (Fig. 1D). This significant lifespan extension occurred in both sexes, with no significant differences between males and females within either condition (Fig. 1E). Because lifespan in *P. dumerilii* is closely linked to sexual maturation, we next asked whether this remarkable increase in longevity was accompanied by a delay in maturation. Highlander worms remained immature throughout the repeated-amputation period and underwent sexual maturation only after amputations ceased (Fig. 1F). The median time to maturation was extended 1.61-fold, from 291 days in Controls to 471 days in Highlander worms. Maturation time did not show sex-biased differences (Fig. S1A–C). We next tested whether maturation time was associated with worm size, estimated from the initial number of segments at the time of collection. In Highlander worms, maturation timing was independent of initial body size, whereas in Controls, smaller worms (40–50 segments) matured significantly later than larger worms (∼60 segments) (Fig. 1G).

Experiment 2 independently reproduced the lifespan extension and delayed maturation observed in Experiment 1. Highlander worms again lived significantly longer than Controls, with a 1.47-fold increase in median lifespan, from 282 days in Controls (n = 16) to 415 days in Highlander worms (n = 16), with no sex-biased differences (Fig. 1H, I). Thus, Control lifespan in both experiments closely matched the culture-wide median of 286 days. Sexual maturation was also delayed 1.47-fold, with median maturation time increasing from 279 days in Controls to 412 days in Highlander worms (Fig. 1J, Fig. S1D–F), and maturation timing was not significantly associated with initial body size (Fig. 1K).

Despite these consistent effects on lifespan and sexual maturation timing, the relationship between regeneration and sexual maturation differed between experiments. In Experiment 1, Highlander worms matured only after repeated amputations had ceased, whereas in Experiment 2, some Highlander worms underwent sexual maturation before completing regeneration, in some cases bearing a total of only 25 segments (Fig. S1G). Thus, although repeated amputation consistently delayed maturation, completion of regeneration was not a prerequisite for sexual maturation.

Together, these experiments demonstrate that repeated amputation and regeneration reproducibly extend lifespan by approximately 1.5-fold-change and substantially delay sexual maturation in *P. dumerilii*, allowing siblings of the same chronological age to follow markedly different developmental trajectories.

### Repeated regeneration delays gametogenesis progression

We next asked whether delayed sexual maturation reflected only a shift in the timing of sexual metamorphosis or whether earlier processes, such as gametogenesis, were also affected. Unlike animals with discrete gonads, *P. dumerilii* develops germ cells within the coelom, where gonial clusters surrounded by sheath cells float freely (Fischer, 1974, 1975). As worms grow, gonial clusters progress through distinct morphological and meiotic stages, providing a means to assess progression of gametogenesis and reproductive development (Kuehn et al., 2022; Ribeiro et al., 2025). To determine whether repeated regeneration affects this progression, in Experiment 3, we compared gonial cluster stages between Highlander worms and their unamputated Control siblings after five months of repeated amputation (5 mpa) (Figs. 2A–F, Figs. S2A–C).

**Figure 2.**
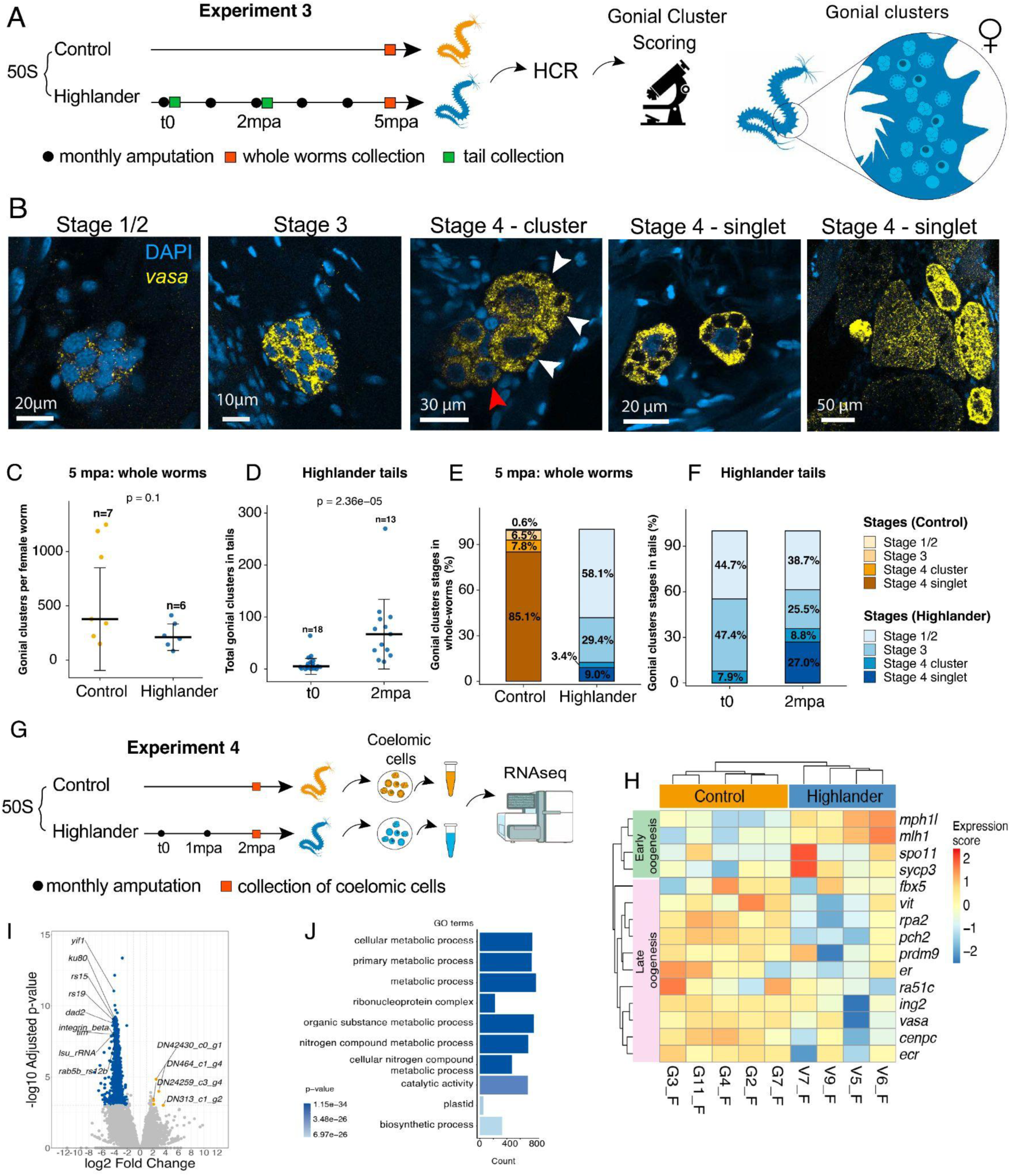
Repeated regeneration delays gametogenesis. (A–F) Experiment 3: morphological assessment of gametogenesis progression. (A) Experimental design. (B) HCR images showing female gonial cluster stages, progressing from early stages (1/2) to Stage 4-singlet. (C) Number of gonial clusters per whole worm at 5 mpa. (D) Number of gonial clusters in amputated tails of Highlander worms, comparing the original tail at t0 with the second regenerated tail at 2 mpa. (E) Distribution of gonial clusters across stages in Control and Highlander worms. (F) Distribution of gonial cluster stages in Highlander tails at t0 and 2 mpa. (G–J) Experiment 4: transcriptomic assessment of gametogenesis progression at 2 mpa. (G) Experimental design. (H) Hierarchically clustered heatmap of selected gametogenesis-related genes. (I) Volcano plot highlighting differentially expressed genes between Control and Highlander siblings at 2 mpa (*p* < 0.001). (J) Top 10 enriched GO terms among genes upregulated in Highlander worms.

To identify and score gonial clusters, we performed whole-mount *in situ* hybridization chain reaction (HCR) targeting *vasa*, a gonial cluster marker, and imaged entire worms and their tails collected after each amputation (Fig. 2A). We adapted the four-stage classification described by Fischer (1974) to account for fixation-induced changes in nuclear morphology in whole-mount HCR (Fig. 2B, Fig. S2A). Because stages 1 and 2 were difficult to distinguish reliably, we combined them into a single category (stage 1/2; Fig. 2B). We also subdivided stage 4 into stage 4-cluster (early stage 4) and stage 4-singlet (advanced stage 4) (Fig. 2B; see Methods for detailed scoring criteria). We focused on females because their gametes exhibit distinct nuclear morphologies that facilitate staging.

By 5 mpa, Highlander worms had undergone five amputation–regeneration cycles, repeatedly regenerating their lost posterior body and remaining approximately at the 50-segment stage (50S; mean = 54, median = 53). In contrast, Control siblings continued growing and reached approximately the 70-segment stage (mean = 74, median = 78; Fig. S2B). At 5 mpa, the average number of gonial clusters did not differ significantly between Highlander and Control worms (Wilcoxon *W* = 33, *p* = 0.1004), although Control worms showed greater variability (Fig. 2C, Fig. S2B). However, these totals combined multicellular gonial clusters with individually counted stage 4-singlets and therefore did not represent equivalent numbers of germ cells across developmental stages. We also compared the original tails removed during the first amputation (t0) with regenerated tails collected after two amputation–regeneration cycles (2 mpa). Despite having comparable numbers of segments (*P* = 0.809, Fig. S2C), tails collected at 2 mpa contained significantly more gonial structures than those collected at t0 (Wilcoxon *W* = 11, *P* < 0.001; Fig. 2D), indicating that gonial clusters continued to accumulate in the posterior body despite repeated amputation and regeneration (Fig. 2D, Fig. S2C).

Although Highlander and Control worms had similar numbers of gonial clusters at 5 mpa, the distribution of gonial cluster stages differed markedly between groups. Stage 4-singlets (the most progressed stage) represented 85.1% of scored structures in Control worms, whereas stage 1/2 clusters (the least progressed stage) accounted for 58.1% in Highlander worms (Fig. 2E). This difference was also evident in the pooled stage distributions (Pearson’s χ²(3) = 3865.8, p < 0.001), although this analysis did not account for clustering of observations within individual worms. Stage-specific comparisons using a negative binomial generalized linear mixed-effects model, including worm identity as a random intercept, identified differences between conditions in stage 1/2 (p < 0.001), stage 4-cluster (p < 0.01), and stage 4-singlet counts (p < 0.001). Together, these observations indicate that Highlander worms retained a greater representation of early gonial cluster stages, whereas Control worms contained predominantly advanced oocytes. In Highlander tails, the pooled stage distributions differed between t0 and 2 mpa (Pearson’s χ²(3) = 17.07, p < 0.001) (Fig. 2F). However, stage-specific comparisons using a negative binomial mixed-effects model did not detect significant differences between time points (all p ≥ 0.73). Thus, although the pooled counts suggested differences in stage composition, the worm-level analysis did not provide sufficient evidence for changes in individual stage counts.

In summary, Experiment 3 shows that repeated amputation does not prevent the accumulation of gonial clusters but is associated with a pronounced delay in their developmental progression, consistent with the delayed sexual maturation observed in Highlander worms.

### Repeated regeneration alters gametogenesis-related gene expression

To investigate how repeated regeneration alters gametogenesis at the gene-expression level, we compared worms at 2 mpa in Experiment 4. At this time point, Control worms had a median of 70 segments, compared with 50 in Highlander worms (Fig. S2F). To assess gametogenesis progression, we collected the coelomic contents, containing the gonial clusters from each worm, and carried out bulk RNA-seq analyses (Fig. 2G). In our previous work, RNA-seq of coelomic contents successfully distinguished both sex-biased transcriptional differences and gametogenesis stage differences among worms between the 40S and 60S developmental stages (Ribeiro et al., 2025). We therefore applied a similar approach here, using our previously assembled reference transcriptome for gametogenesis progression.

We then generated a hierarchically clustered heatmap of 15 selected genes to compare the expression patterns of meiosis-and oogenesis-related genes in RNA-seq data from the coelomic contents of Highlander and Control worms, regardless of whether they met the stringent *P* < 0.001 cutoff (Fig. 2H). These genes were selected based on meiosis-associated GO terms identified in our previous study (Ribeiro et al., 2025), along with *vasa*, *ecr*, *er*, and *vitellogenin*, which increase in expression during gametogenesis (Fig. S2D). The heatmap separated Highlander and Control samples and revealed two groups of genes associated with early and late oogenesis (Fig. 2H). Notably, late-oogenesis genes showed higher expression in Control worms. Among the selected genes, only *mph1l*, a gene associated with early oogenesis, was significantly differentially expressed between Highlander and Control worms (*P* < 0.05; Fig. S2E). Thus, the expression patterns of oogenesis-related genes indicate that, by only 2 mpa, Control worms had progressed further through gametogenesis than Highlander worms.

Additionally, our differential expression analysis with a cutoff of *p* < 0.001 identified 1,853 genes up-regulated in Highlander coelomic contents and only 6 genes up-regulated in Controls (Fig. 2I; Fig. S2G–H). While all 6 transcripts up-regulated in Controls lacked functional annotation, many of the genes up-regulated in Highlanders were annotated, allowing us to perform gene ontology (GO) enrichment analysis. The top GO terms among genes up-regulated in the Highlander group were mainly associated with metabolic processes, which accounted for 6 of the top 10 GO terms (Fig. 2J).

Together, these results show that repeated regeneration is associated with delayed gametogenesis and a transcriptional profile characterized by increased expression of metabolism-related genes.

### Repeated regeneration delays somatic pigmentation

We next tested whether repeated regeneration affected development beyond reproductive progression. In *P. dumerilii*, segment number is a useful morphological proxy for developmental stages. As worms grow, germ cells expand and begin to differentiate, and once worms reach a threshold of approximately 70 segments, they become predisposed to sexual maturation. Thus, worms with more segments are generally more advanced developmentally and closer to maturation and death. However, because amputation removes segments, segment number cannot be used directly to compare overall developmental progression between Highlander and Control worms undergoing repeated regeneration.

We therefore examined two additional morphological markers that could be used to compare developmental progression in *P. dumerilii*: the number of pigment spots, or chromatophores, and eye size (Fig. 3A–J; Fig. S3A–O). In unamputated worms, both traits increased with developmental stage, supporting their use as markers of somatic growth in *P. dumerilii* (Fig. 3A; Fig. S3E). We expected that, if Highlander worms remained phenotypically younger, eye size and pigment number would either remain unchanged or increase less than in their Control siblings.

**Figure 3.**
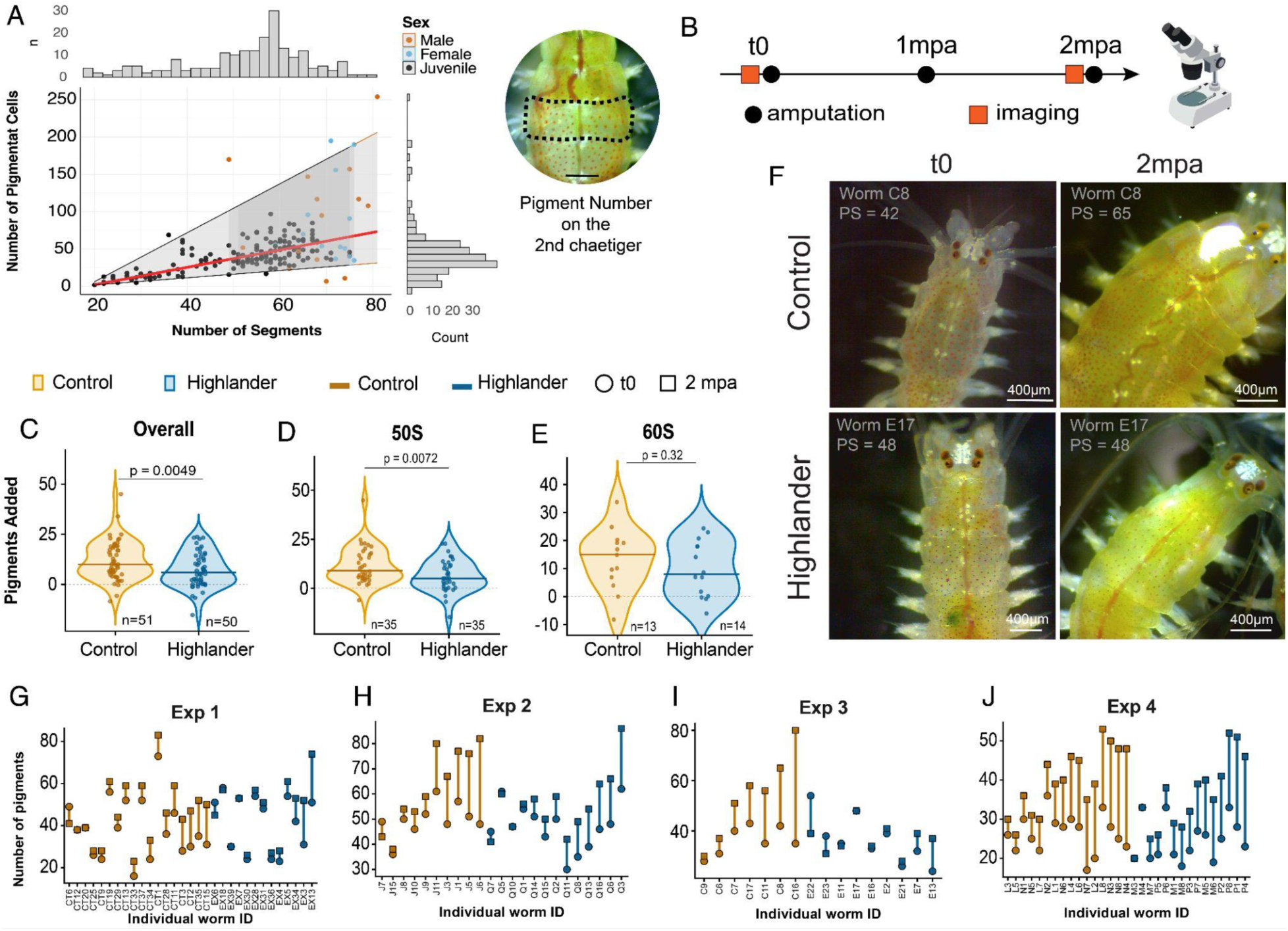
Repeated regeneration delays pigment addition. (A) Number of pigment spots on the second chaetiger across developmental stages (left) and light microscopy image of the dorsal region of the second chaetiger (right); scale bar, 400 μm. (B) Experimental design used to assess differences in pigment addition. (C–E) Number of pigment spots added from t0 to 2 mpa, showing the overall distribution (C), 50S worms only (D), and 60S worms only (E). Violin plots show individual animals, group medians, and a zero-change reference line. (F) Light microscopy images of the anterior body of *P. dumerilii*, showing pigmentation in dorsal view. (G–J) Changes in the number of pigment spots for individual worms in Experiments 1–4.

First, we quantified pigment addition in Highlander and Control siblings before amputation and at 2 mpa (Fig. 3B) (See Methods). At 2 mpa, Highlander worms added fewer pigment spots than Control siblings in the overall analysis, which combined longitudinal data from four independent experiments. The mean number of pigments increased from 35.8 to 51.5 in Control worms, but only from 35.5 to 36.3 in Highlander worms. Thus, over 2 mpa, Control worms added approximately 15 more pigments than Highlander worms, according to our linear (mixed) model analyses (β = −0.099 ± 0.035, SE = 0.035, test statistic = −2.81, p = 0.0049; Fig. 3C).

Our initial analysis included Control and Highlander siblings from different experiments that varied in their initial developmental stage, as determined by segment number. To assess whether developmental stage influenced pigment addition, we analyzed 50S and 60S worms separately. Among 50S worms, Controls added significantly more pigment than Highlanders (*p* < 0.001; Fig. 3D). A similar trend was observed among 60S worms, although the difference was not statistically significant (*p* = 0.32; Fig. 3E). These results also support delayed pigment accumulation in Highlander worms, particularly at the 50S stage.

We next verified whether this pattern was reproducible across independent experiments. Analysis of individual worms further showed that Control worms generally gained more pigments over 2 months, whereas Highlander worms showed little or no increase (Fig. 3G–J). On average, pigment spot addition was consistently reduced in Highlander worms across Experiments 1–4, although only Experiment 1 showed statistical difference (Fig. S3A–D). Together, these findings suggest that repeated regeneration slows the addition of pigments in the anterior body.

We next examined eye growth, measured as interocular distance (IOD; see Methods). Eye size tends to increase during regular development. If repeated regeneration delayed developmental somatic growth, we expected IOD to increase less in Highlander worms than in their Control siblings. In contrast with our expectations, IOD changed significantly in both Control and Highlander worms at 2 mpa and 5 mpa in the overall analyses (Fig. S3F, G). However, the pattern varied among experiments. In Experiments 1 and 2, IOD changed significantly in both conditions, whereas Experiment 4 showed no significant change in either condition (Fig. S3H, I, K). Only Experiment 3 showed the pattern predicted by our hypothesis: IOD changed significantly in Control worms (p < 0.001), while Highlander worms remained close to a fold-change of 1 (p = 0.63; Fig. S3J). Across experiments, IOD fold-change in Control worms tended to deviate more from 1, where a value of 1 indicates no change in IOD over time, whereas Highlander worms tended to remain closer to 1 (Fig. S3H–O). However, this difference between conditions was not consistently supported statistically across experiments. Thus, with our current approach, we cannot conclude that repeated regeneration delays changes in eye size.

Overall, changes in pigment number indicate that repeated regeneration is associated with slower somatic growth in regions of the body that are not directly regenerated. These results suggest that the effects of repeated regeneration extend beyond reproductive progression, and are also associated with changes in somatic developmental progression.

### Individuals maintain regenerative competence across repeated amputations

We next examined whether individuals undergoing repeated regeneration maintain their regenerative competence over time. Using the same sibling cohort from the lifespan experiment (Experiment 2; Fig. 1G–J), we assessed regenerative competence by counting the number of segments regenerated between successive monthly amputations (Fig. 4A). Because amputations occurred at approximately 30-day intervals, this approach allowed us to compare the extent of posterior regeneration over similar periods across successive cycles.

**Figure 4.**
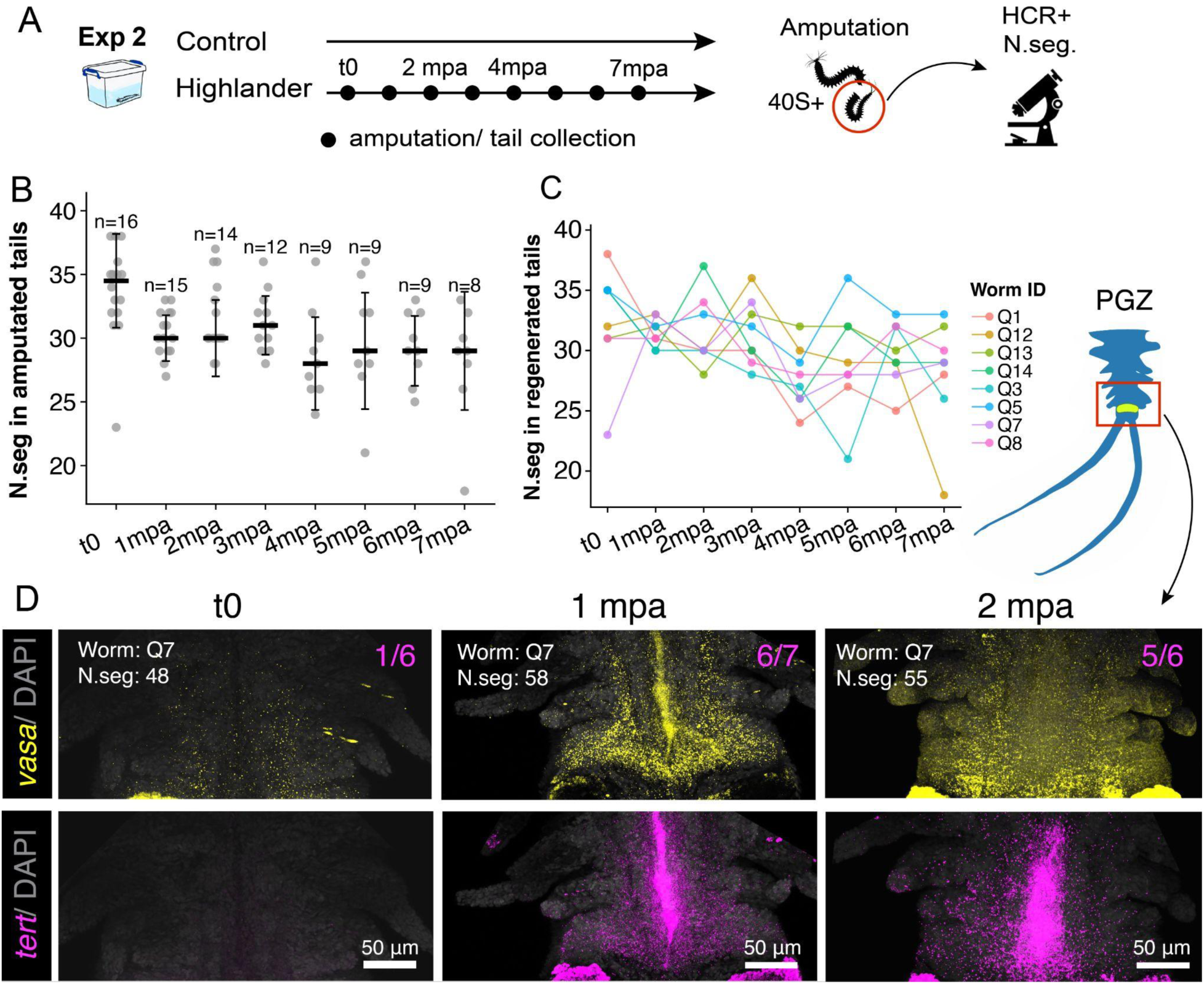
Regenerative capacity remains robust over repeated amputation-regeneration cycles. (A) Experimental design (Experiment 2). The amputation site was kept constant across cycles. Tails were collected at each amputation, and their segments were counted. t0 corresponds to the first amputation of sibling worms initially at various developmental stages (40S or later). Worms underwent a maximum of seven amputations. Tails collected at t0, 1 mpa, and 2 mpa were also fixed for HCR after segment counting. (B) Number of segments in amputated tails across amputation cycles. (C) Number of segments in amputated tails from worms that survived all seven amputations and subsequently matured. (D) HCR detection of *tert* expression in the posterior growth zone (PGZ) in tails collected longitudinally from the same worm (ID: Q7); *vasa* was used as a positive control. Fractions indicate the number of *tert*-positive worms relative to the total number examined.

At the beginning of the experiment, before amputations started, Highlander worms ranged from 48 to 72 segments, representing developmental stages from the 40S onward. Among Highlander worms that remained juvenile, the mean number of regenerated segments remained similar across seven amputation–regeneration cycles, ranging from 31.7 segments at 2 mpa to 28.1 segments at 7 mpa. We detected no significant differences in the number of regenerated segments among cycles from 1 to 7 mpa (all *p* ≥ 0.069; Fig. 4B, C; Fig. S4B, C).

We next tested whether the initial developmental stage affected regenerative output. Worms generally regenerated approximately half of their body, and the proportion regenerated was not correlated with their initial number of segments (Spearman’s ρ = −0.24, p = 0.39; Fig. 4A, B; Figs. S4B–C). Thus, within the developmental range examined, regenerative output was largely independent of the initial developmental stage, with worms regenerating at a rate of approximately one segment per day.

Because regenerative competence may rely strongly on mechanisms of DNA maintenance, we examined the posterior growth zone (PGZ) in regenerated tails and compared it with the original tail collected at t0, before the first amputation. Using HCR, we analyzed the expression of *tert (telomerase reverse transcriptase)*, which encodes the catalytic subunit of telomerase, an enzyme involved in telomere maintenance. *vasa*, which is known to be expressed in the PGZ, was used as a positive control. Detectable *tert* expression in the PGZ was uncommon at t0, observed in only 1 out of 6 worms, but was present in 6 out of 7 worms at 1 mpa and 5 out of 6 worms at 2 mpa (Fig. 4C; Fig. S4D, D). These observations indicate that *tert* expression detectable by HCR *in situ* in the PGZ is more frequent following regeneration and is maintained across at least two repeated regeneration cycles.

### Amputation induces developmental-stage-associated genes in the head

Regeneration and reproduction in *P. dumerilii* are closely associated with neuroendocrine regulation, especially via hormones released from the brain (Hauenschild, 1955, 1960, 1966). We therefore asked whether amputation and regeneration induce changes in the head that could preserve a younger physiological state. We first characterized transcriptional changes in the head associated with developmental stages without any amputations. Heads were collected for RNA sequencing (RNA-seq) from individuals originating from independent batches and representing the 40S, 50S, and 60S developmental stages (Fig. 5A). Quality-control analyses revealed substantially greater variability in the mapping profiles of the 50S samples compared with the 40S and 60S samples (Fig. S5A–E). We therefore excluded the 50S group from the final analysis. The subsequent analysis compared 40S and 60S animals, with six biological replicates retained for each group. Sample-to-sample correlation analysis further confirmed the relationships among the retained biological replicates (Fig. 5B, Fig. S5F).

**Figure 5.**
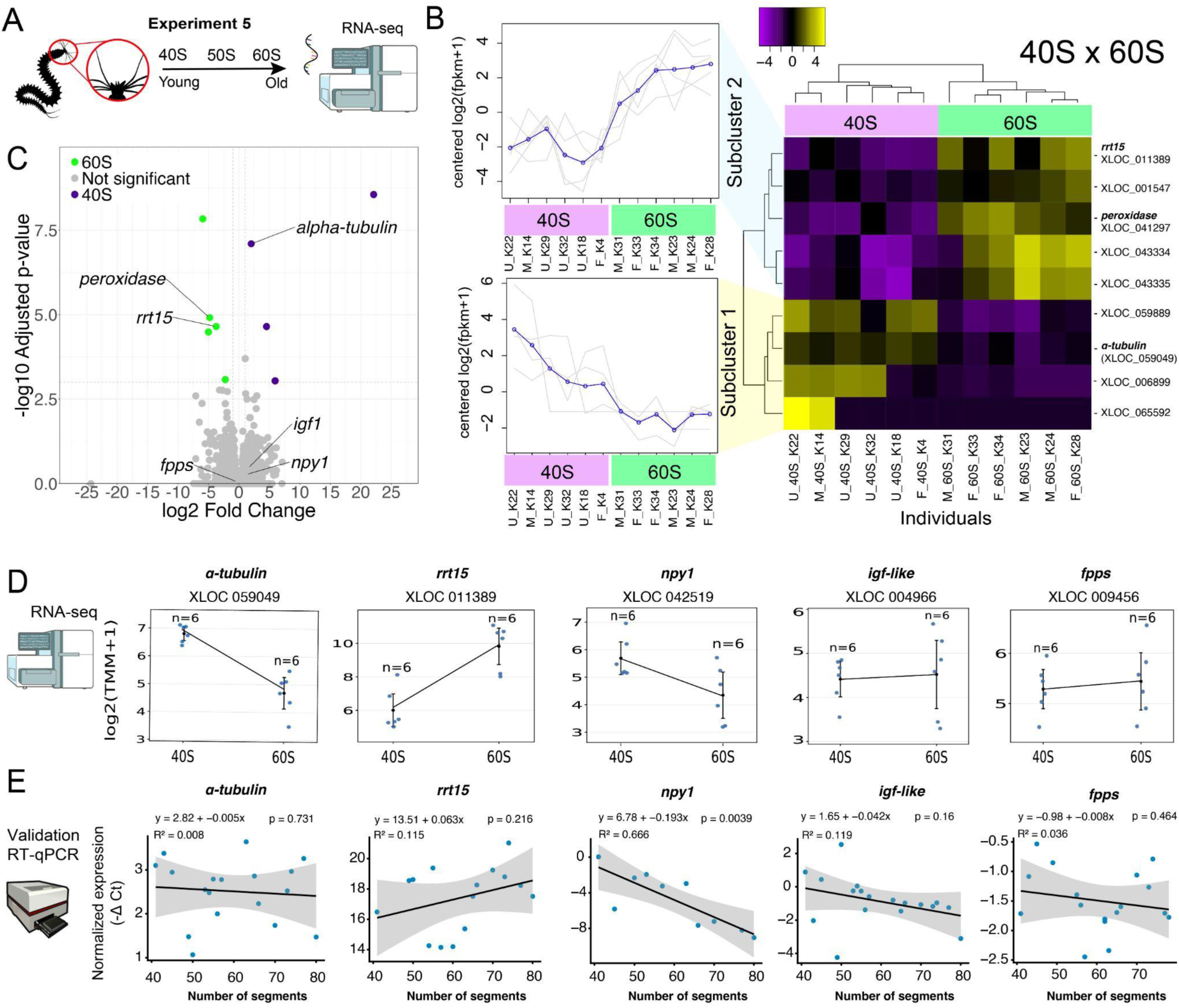
Transcriptomic identification and validation of candidate markers associated with developmental age in the *Platynereis* head. (A) Experimental design for RNA-seq analysis of heads collected from juveniles spanning developmental stages of 40–60 segments (40S–60S). (B) Heatmap of differentially expressed genes (DEGs; *p* < 0.001), with hierarchical clustering revealing distinct gene-expression subclusters. (C) Volcano plot highlighting differentially expressed genes in purple and green, together with selected genes of interest that did not meet the differential-expression threshold. (D) RNA-seq expression profiles of selected candidate genes potentially involved in brain-mediated regulation of regeneration. (E) RT-qPCR validation of selected candidate genes across individuals spanning different developmental stages.

Differential-expression analysis identified nine genes that differed significantly between 40S and 60S heads using a significance threshold of *p* < 0.001 (Fig. 5B, C). Clustering of these genes revealed two major expression subclusters. Genes in Subcluster 1 showed higher expression in younger 40S individuals, whereas genes in Subcluster 2 showed higher expression in older 60S individuals (Fig. 5B). We next examined the annotations of these nine differentially expressed genes. Among genes with annotations corresponding to known proteins, *α-tubulin* showed higher expression in 40S animals, whereas *peroxidase* and *rrt15* (regulator of rDNA transcription protein 15) showed higher expression in 60S animals.

In addition to these differentially expressed genes, we examined a broader set of candidate genes that did not meet the statistical threshold for differential expression but were selected based on known or proposed roles in neuroendocrine signaling, regeneration, growth, reproduction and development. We focused on genes associated with methyl farnesoate (MF), GnRH, IGF, and NPY signaling pathways, which have been implicated in growth and regeneration, reproductive regulation, growth and metabolism, and stress responses, respectively (Álvarez-Campos et al., 2023; Andreatta et al., 2020; Choi et al., 2025; Reichmann & Holzer, 2016; Schenk et al., 2016). Genes identified through our gene ontology annotations as associated with MF signaling did not show clear clustering by developmental stage (Fig. S5G–I). Similarly, genes associated with GnRH signaling showed no consistent age-associated pattern (Fig. S5J). In contrast, genes associated with IGF and NPY signaling pathways showed expression patterns that clustered the younger 40S heads together (Fig. S6J). Based on these patterns, we selected *npy1* and *igf-like* for RT-qPCR validation, together with *α-tubulin* and *rrt15*, which were significantly differentially expressed between 40S and 60S heads. We also selected *fpps* (farnesyl diphosphate synthase) for validation. Although *fpps* did not show differences across developmental stages, it was included because of its involvement in the methyl farnesoate biosynthetic pathway (Li et al., 2025).

Selected candidate genes were subsequently examined by RT-qPCR across individuals spanning different developmental stages, allowing their expression patterns to be compared with those observed by RNA-seq (Fig. 5D, E). The expression patterns of rrt15, npy1, and igf-like were broadly consistent between the two approaches. npy1 expression significantly decreased with developmental stage (slope = −0.193, p < 0.01), whereas rrt15 showed a non-significant trend toward increased expression with developmental stage (slope = 0.063, p > 0.05). igf-like also showed a tendency toward decreased expression, although this relationship was not statistically significant (slope = −0.042, p > 0.05). In contrast, α-tubulin, which was among the most strongly differentially expressed genes in the RNA-seq analysis and showed lower expression in 60S than in 40S animals, did not show an age-associated trend by RT-qPCR (slope = −0.005, p > 0.05). Similarly, fpps expression remained stable across developmental stages, with no significant relationship between expression and developmental stage detected by regression analysis (slope = −0.008, p > 0.05).

Having determined molecular markers that may represent younger (40S) vs older (60S) worm heads, we next asked whether amputation increases the expression of younger head-associated genes in more developmentally advanced worms (50S). To do this, we amputated 50S worms in half (between segment 25-26) and collected their heads 2 hours after amputation (2 hpa) (Fig. 6A). RT-qPCR analysis showed that, at 2hpa, *α-tubulin* (p<0.05, Fig. 6B) and *igf-like* (p<0.01, Fig. 6C) increases significantly. No significant changes in the expression of *rrt15*, *npy1*, or *fpps* were detected at 2 hpa (Fig. 6D–F).

**Figure 6.**
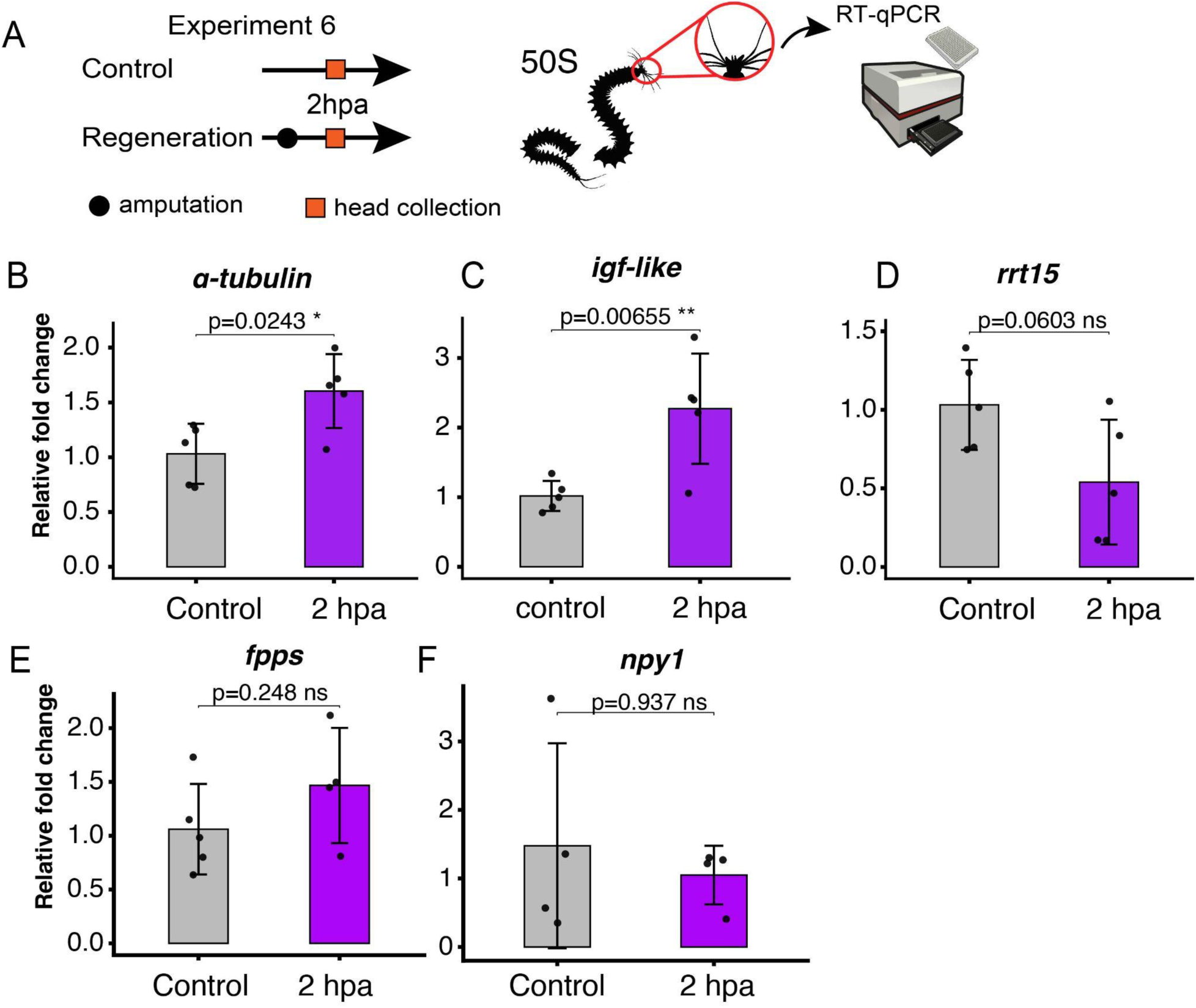
Head gene-expression responses following posterior amputation. (A) Experimental design used to assess changes in head gene expression after amputation. 50S worms were amputated at mid-body, removing approximately 25–30 segments. (B–F) RT-qPCR analysis of *α-tubulin* (B), *igf-like* (C), *rrt15* (D), *fpps* (E), and *npy1* (F) expression following amputation.

Together, the RNA-seq and RT-qPCR analyses identified candidate genes with expression patterns associated with younger and older developmental stages in the juvenile head. RNA-seq identified differentially expressed genes, including *α-tubulin* and *rrt15*, whereas RT-qPCR revealed a significant decrease in *npy1* expression with developmental progression. We next investigated whether these developmental-stage-associated genes also responded acutely to injury. Although *α-tubulin* and *igf-like* did not show significant developmental-stage-associated changes by RT-qPCR, both were significantly upregulated in the head at 2 hpa. These findings suggest a partial overlap between transcriptional programs associated with juvenile development and the acute injury response in the head, identifying potential candidates for further investigation of the molecular mechanisms linking injury, regeneration, and developmental progression.

## Discussion

Our findings identify a natural phenotype in which repeated regeneration delays the terminal reproductive transition and extends lifespan in the semelparous annelid *Platynereis dumerilii*. We refer to this phenotype as the Highlander effect. Following repeated removal of the posterior body, animals remained in a juvenile developmental condition longer than age-matched sibling controls, progressed more slowly through gametogenesis and sexual metamorphosis, accumulated pigmentation more slowly, retained regenerative competence, and lived approximately 1.5-fold longer. These findings reveal that repeated regeneration upon amputation promotes systemic effects, i.e. changes in organism-wide physiology.

### The systemic effects of regeneration on developmental trajectories

Tissue injury triggers physiological responses that extend beyond the damaged tissue, including systemic inflammation, endocrine signaling, and metabolic adjustments associated with repair (Rennolds & Bely, 2023). These responses can influence cellular activity in distant tissues, as well as organismal growth and reproductive physiology. For instance, limb amputation triggers cell-cycle entry in the uninjured contralateral limb in axolotls (Johnson et al., 2018). In lizards, injury can also alter reproductive investment, as tail autotomy followed by regeneration increases egg and hatchling size (Beatty et al., 2021). Beyond these systemic responses, injury and regeneration can induce more profound changes in developmental and physiological states of some invertebrates. For example, the ctenophore *Mnemiopsis leidyi* can undergo reverse development from the mature form to larval stages upon injury (Soto-Angel & Burkhardt, 2024). Similarly, the cnidarian *Turritopsis dohrnii* can revert from the medusa to the polyp stage in response to injury, starvation, environmental stress, or senescence (Piraino et al., 1996).

Regeneration in aged sexual planarians has been shown to reverse age-associated cellular, molecular, and physiological changes in tissues both near and far from the amputation plane (Dai et al., 2025). Repeated amputation-regeneration in the asexual annelid *Paranais litoralis* resulted in rejuvenation of the somatic cells and lifespan extension (Martínez, 1996). Collectively, these studies illustrate how injury and regeneration can influence physiological processes throughout the organism, with consequences for development, reproduction, and aging.

In our experiments, each amputation removed approximately half of the body in *P. dumerilii*, and restoration of this tissue required approximately one month. Each cycle therefore imposes several regeneration steps during this period: injury response, wound healing, blastema formation, blastema differentiation, re-establishment of the posterior stem cell zone (PGZ), and recovery of body size. The repeated amputation regime maintained animals in recurrent cycles of injury, regeneration, and compensatory growth. A striking systemic effect of this repeated regeneration was the delay in reproductive progression and maintenance of young juvenile characteristics. Control females progressed further through gametogenesis than Highlander females, as evidenced by gonial cluster morphology and the expression of oogenesis-associated genes, indicating differences in reproductive development between siblings of the same chronological age. Highlander worms also exhibited slower pigment accumulation and tended to remain smaller, suggesting that the effects of repeated amputation extend beyond reproductive tissues. These findings indicate that repeated regeneration is associated with delayed reproductive maturation and changes in somatic development, consistent with a broader influence of regeneration on developmental progression.

### Retention of regenerative competence across repeated amputation-regeneration cycles

Regenerative capacity usually declines with age across animals (Yun, 2015). For example, in the short-lived killifish *Nothobranchius furzeri*, caudal fin regeneration progressively declines with age (Wendler et al., 2015). In addition, the ability to regenerate can fluctuate during development, with periods of permissive or non-permissive regenerative capacity. For instance, in *Xenopus* tadpoles, tail regeneration is initially robust, transiently declines during a refractory period, and is subsequently restored before being largely lost during metamorphosis (Kakebeen & Wills, 2019; Williams et al., 2021). These findings illustrate that regenerative capacity is influenced by both organismal aging and developmental state.

In *P. dumerilii*, regenerative capacity progressively declines as juveniles approach sexual maturation, with increasing rates of regeneration failure and ultimately a complete loss of regenerative ability upon reaching sexual maturity (Álvarez-Campos et al., 2023; Hauenschild, 1966; Metzger & Özpolat, 2024). We found that Highlander worms retained regenerative competence across time, supporting the idea that these worms retained a younger juvenile state, compared to their siblings that matured sexually and lost the ability to regenerate. In *P. dumerilii,* regenerative outcomes are often associated with a brain hormone that promotes somatic growth and regeneration while suppressing sexual maturation (Hauenschild, 1956, 1960, 1966, 1974). The sesquiterpenoid hormone methyl farnesoate has been identified as an important component of this brain-derived activity (Schenk et al., 2016). Decapitation, which effectively removes the brain – the source of the hormone – accelerates maturation and abolishes regenerative competence (Hauenschild, 1956, 1960). In contrast, transplantation of a younger juvenile’s brain into a decapitated worm can restore growth and regeneration and inhibit reproductive progression (Hofmann, 1976). Retention of regenerative competence in Highlander worms may therefore be both a component and a consequence of the extended juvenile neuroendocrine state: the brain potentially regulates regenerative competence, but injury-induced systemic responses may in turn act on the brain to modify transcriptional programs and increase the production of this proregenerative hormone. Together, these findings suggest that repeated regeneration uncouples chronological age from reproductive and somatic developmental progression.

In addition, in some organisms, repeated amputation-regeneration results in progressive failure and loss of fidelity of regeneration. For example, in axolotls, repeated limb amputation leads to declining regenerative fidelity and eventual regeneration failure, associated with dysregulated amphiregulin expression (Bryant et al., 2017). In contrast, Highlander worms retained regenerative competence across repeated amputation cycles despite repeated tissue loss, without an apparent decline in posterior growth rates. The sustained expression of *tert* and *vasa* in the posterior growth zone (PGZ) may provide insights into the cellular mechanisms underlying this regenerative resilience in *P. dumerilii*. Telomerase activity supports telomere maintenance during regeneration in zebrafish and asexual planarians (Anchelin et al., 2011; Dai et al., 2025). Loss of *tert* impairs cardiac regeneration in zebrafish and tert RNAi reduces regeneration speed in planarians (Bednarek et al., 2015; Dai et al., 2025). Similarly, *vasa* is associated with proliferative stem and progenitor cell populations and is expressed during posterior regeneration in *P. dumerilii* (Gazave et al., 2013; Kozin & Kostyuchenko, 2015; Planques et al., 2019). Therefore, sustained *tert* and *vasa* expression may reflect the continued maintenance of regeneration-associated cellular programs in Highlander worms. Whether these programs actively preserve regenerative competence or are maintained as a consequence of delayed developmental progression remains to be determined.

### Short and long-term hormonal changes induced by injury and regeneration

Injury can elicit short-term (immediate) systemic changes and these quick responses can, in turn, lead to long-lasting effects in the organism. We therefore examined transcriptional changes in brain hormonal pathways, particularly methyl farnesoate (MF) signaling. Previous work showed that MF concentrations in *P. dumerilii* heads decline by approximately 50% between premature and sexually mature animals, with reduced expression of its putative receptor met in eleocytes (Schenk et al 2016). A subsequent study identified extensive transcriptomic and proteomic changes in worm heads during the transition from premature to mature stages (Schenk et al., 2019). In our comparison of younger and older, unamputated juveniles, MF-pathway genes showed no consistent age-dependent expression differences (Fig. S5G–I). We postulate that MF pathways may change in the long term, as this hormone is known to promote regeneration and growth, while its marked decline during sexual maturation is associated with extensive metabolic changes (Hauenschild, 1966; Hofmann, 1976; Schenk et al., 2016, 2019). Our results indicate that the transcriptional changes associated with later maturation are not necessarily evident during earlier juvenile development. However, this does not rule out regulation of MF signaling at the hormonal level.

We also identified other candidate components of the neuroendocrine state associated with regeneration. We first examined molecular changes occurring in the head during juvenile development to identify candidate genes and then investigated whether their expression levels changed in the short-term after injury. We found that the expression of *npy1*, *α-tubulin*, and *igf-like* tended to decrease with developmental progression, whereas *rrt15* expression increased. We then found that the expression of both *α-tubulin* and *igf-like* rapidly increased in expression following posterior amputation. Thus, the acute injury response partially mimics the transcriptional program of the young juvenile heads. This overlap does not demonstrate that injury rejuvenates the head, but that the removal of the posterior of the body reactivates components of a molecular program associated with younger animals. Determining whether this response is transient or sustained in the long term, and across amputation-regeneration cycles will be necessary to establish whether repeated amputation merely triggers successive acute responses or maintains a chronically altered neuroendocrine state.

Among neuroendocrine candidates, insulin/IGF signaling provides a particularly plausible connection between injury, systemic physiology, and developmental timing. Across diverse animals, this pathway coordinates growth, metabolism, reproduction, stress responses, and lifespan, although its effects are strongly context-dependent (Clemmons, 2012). In *Drosophila,* damage to imaginal discs induces secretion of DILP8, an insulin-like peptide that delays developmental progression and extends the larval stage (Colombani et al., 2012; Garelli et al., 2015). Similarly, insulin/IGF signaling could link repeated injury to delayed maturation and the metabolic demands of regeneration in *P. dumerilii*. Consistent with altered systemic physiology, Highlander worms showed enrichment of metabolic pathways in their coelomic contents (Fig. 2J).

These contents include eleocytes, cells involved in nutrient storage and metabolism, and gonial clusters containing developing germ cells (Fischer, 1974, 1975; Meisel, 1990; Schenk et al., 2016; Schröer, 2004). The metabolic shift detected in coelomic contents in Highlanders may reflect changes in eleocyte metabolic activity, differences in the relative abundance of eleocytes and gonial clusters, or both. Although bulk transcriptional analyses cannot distinguish these possibilities, metabolic enrichment identifies coleomic physiology as a potential component of the relationship between repeated regeneration and delayed sexual maturation in Highlander worms.

### Regeneration at the intersection of reproduction and aging

The delay in reproduction places the Highlander effect within the broader relationship among reproduction, somatic maintenance, and lifespan. In P. dumerilii, sexual maturation culminates in a single reproductive event followed by death, making the terminal reproductive maturation a major determinant of lifespan. Recent work demonstrated that loss of the light-receptive cryptochrome L-Cry extends lifespan by approximately 1.40–1.45-fold change through prolonged juvenile development and delayed sexual maturation, as shown by gametogenesis progression (Andreatta et al., 2026). At the neuroendocrine level, L-Cry targets the atypical nuclear receptor *nr0b1/2* (orthologous to DAX-1/SHP), shifting the expression of conserved steroidogenic enzymes, estrogen/nuclear receptors, and insulin/IGF signaling components alongside the longevity factor *pdp1* (Andreatta et al., 2026). While it remains unknown whether L-Cry mutants and Highlander worms share identical downstream mechanisms, these signatures demonstrate that light perception shapes lifespan through ancient hormonal networks.

The relationship between reproduction and lifespan is well established across animals. In *Caenorhabditis elegans*, germline ablation extends lifespan through mechanisms involving insulin/IGF signaling and nuclear hormone receptors (Hsin & Kenyon, 1999). Similarly, reproductive investment is associated with survival costs in *Drosophila*, although these trade-offs do not necessarily result from competition for limited resources (Flatt, 2011). Classical theories of aging suggest two nonexclusive explanations for the Highlander effect. First, according to the disposable soma theory, delaying reproduction could redirect limited resources from germline development toward regeneration, cellular repair, and somatic maintenance, thereby reducing the later-life costs associated with reproductive investment (Kirkwood, 1977). Second, aging is characterized by a progressive loss of physiological integrity, driven by interconnected molecular and cellular hallmarks, that leads to impaired function and increased vulnerability to death (López-Otín et al., 2013, 2023). The retention of regenerative competence, sustained expression of regeneration-associated genes, and altered metabolic profiles observed in Highlander worms are consistent with these possibilities.

Importantly, because sexual maturation is followed by death in *P. dumerilii*, extended lifespan does not necessarily indicate slower biological aging. Repeated regeneration may actively stimulate repair programs that preserve physiological integrity and slow the accumulation of age-related damage. Alternatively, it may primarily postpone the transition to sexual maturity, independently of changes in rate of physiological decline. Distinguishing these possibilities will require measurements of age-associated molecular and physiological changes independently of reproductive development. Thus, the Highlander effect provides an opportunity to investigate whether regeneration extends lifespan solely by delaying reproductive maturation or also by modifying the underlying processes of biological aging.

### Conclusions and future directions

In conclusion, repeated regeneration maintains *P. dumerilii* in a juvenile, regeneration-competent state and delays its terminal reproductive transition. We term this systemic and organism-wide response the Highlander effect and propose that tissue loss repeatedly resets or sustains neuroendocrine programs that favor regeneration over reproduction. The central implication of the Highlander effect is that repeated regeneration uncouples chronological age from developmental progression and raises the possibility that regenerative demand can also alter the trajectory of biological aging.

Key questions now concern the signals that trigger and maintain this state. Because each amputation removed approximately half of the body, it remains unknown whether the Highlander effect scales with injury magnitude or can also be induced by smaller tissue loss. Sustained expression of *tert* and *vasa* may contribute to regenerative maintenance or reflect the persistence of regeneration-associated cell populations. Posterior amputation also alters head gene expression within two hours, including increased *igf*-*like* and α-tubulin expression, suggesting rapid communication between local injury and distant regulatory tissues. In this context, the insulin/IGF pathway is a particularly compelling candidate for connecting injury sensing with the systemic state of Highlander animals, especially given the metabolic differences observed in their coelomic physiology. Testing the causal role of insulin/IGF signaling will determine whether this pathway contributes to the maintenance of regenerative competence, altered metabolism, delayed gametogenesis, or multiple components of the phenotype. In parallel, combining eleocyte and gonial-cluster quantification with cell-type-resolved transcriptomics and measurements of vitellogenin will reveal whether the metabolic state of Highlander worms results from changes in coelomic cell composition, altered eleocyte physiology, or both. Methylfarnesoate (MF) signaling also remains a leading candidate mechanism linking regeneration and reproduction. Direct measurements of MF levels, together with functional manipulation of the pathway, will therefore be essential for determining whether repeated amputation maintains the neuroendocrine state that supports regeneration while delaying sexual maturation. Finally, we establish *P. dumerilii* as a powerful system for distinguishing resource-allocation trade-offs from active maintenance programs and for testing how regeneration reshapes the relationship between development, reproduction, and aging.

### Limitations of this study

Although our findings establish the Highlander effect, several limitations should be considered when interpreting these results. First, because sexual maturation, reproduction, and death are tightly coupled in semelparous *P. dumerilii*, lifespan extension cannot by itself distinguish delayed maturation from slower biological aging. Second, pigmentation reflects somatic developmental progression in juveniles but is not an independent biomarker of aging, particularly because pigmentation decreases during sexual maturation. Third, gametogenesis was examined primarily in females due to the gradual, morphologically distinguishable stages of oogenesis. In males, the rapid onset of spermatogenesis may have limited our ability to detect regeneration-dependent effects. Finally, our experimental design does not distinguish the contributions of injury, tissue loss, and regenerative growth to the Highlander effect. Future studies using sham injuries, graded amputations, varied amputation frequencies, and inhibition of regeneration will help determine the mechanisms driving this phenotype and its relationship to regenerative demand.

## Acknowledgements

This research was funded through NIH MIRA R35GM138008 (PI: DBÖ) and Junior Innovator Award from the Center for Regenerative Medicine at Washington University (PI: RPR). We thank the Genome Technology Access Center at the McDonnell Genome Institute at Washington University School of Medicine for help with genomic analysis. The Center is partially supported by NCI Cancer Center Support Grant #P30 CA91842 to the Siteman Cancer Center from the National Center for Research Resources (NCRR), a component of the National Institutes of Health (NIH), and NIH Roadmap for Medical Research. This publication is solely the responsibility of the authors and does not necessarily represent the official view of NCRR or NIH.

## Methods

### Chronological age, developmental stage, and aging in Platynereis dumerilii

Throughout this study, we distinguish chronological age from developmental stage. Chronological age refers to the time elapsed since birth, measured in days, whereas developmental stage refers to the position of an individual along the juvenile growth and maturation trajectory. In *P. dumerilii*, the number of segments provides a useful morphological proxy for developmental stage during juvenile growth, and we therefore refer to animals as 40S, 50S, or 60S according to their number of segments. We use physiological state more broadly to describe the combined reproductive, somatic, regenerative, and molecular condition of the animal. We use these measurements to investigate whether repeated regeneration alters the relationship between chronological age and developmental progression, and whether this altered trajectory is accompanied by broader physiological changes associated with biological aging.

### Regeneration experiments

#### Assessment of Lifespan Duration: regeneration Experiments 1 and 2

To compare lifespan duration between repeatedly amputated worms (Highlanders) and their uncut control siblings, we collected batches of sibling worms derived from single-pair matings 181 days after birth. Two independent batches were used and analyzed separately as well as in combination. All worms were maintained in individual cultures, with sizes ranging from 36 to 68 segments at the start of the experiment (Fig. 1F, J).

The two batches were maintained under different housing conditions. Worms from the August 18, 2023, fertilization batch (Experiment 1) were housed in 50 mL round containers. Some worms constructed and inhabited new tubes along the ridges near the upper edges of the containers, where they were exposed to desiccation. Mortality before maturation was 23.6% in Control worms and 55% in Highlander worms, resulting from desiccation or unknown causes (Suppl. Figs. 1B, C). To reduce this source of mortality, we performed Experiment 2 with worms from January 29, 2025, using 50 mL square containers without ridges (Fig. 1G–J; Suppl. Fig. 1D–F). Under these conditions, no worms died before maturation.

The amputation site was determined based on worm size, measured by the number of segments. Worms with 50 or fewer segments were amputated between segments 25 and 26, leaving an anterior fragment containing 25 segments. Worms with 60 segments or more were amputated between segments 32 and 33, leaving an anterior fragment containing 32 segments. Repeated amputations were performed at the same site every month, after an average of approximately 30 days of regeneration.

#### Assessment of Gametogenesis Progression: Regeneration Experiment 3

We selected worms with 50 segments (50S) in accordance with previous research showing lifespan extension after a single amputation at this stage. In those studies, both 50S and 60S worms that were amputated and regenerated once exhibited extended lifespan. However, because 60S worms are already advanced in gametogenesis, we selected 50S worms to enable a clearer comparison of gametogenesis progression. At the 50S stage, gonial clusters are scattered and include both mitotic and meiotic stages, whereas in 60S worms they are predominantly meiotic. This allows comparison of progression from mitotic to meiotic stages across a broader range. At the 50S stage, sex identification based on gonial cluster morphology and RT-qPCR is highly reliable. In contrast, 40S worms often exhibit gonial clusters at very early stages that cannot be accurately sexed. Thus, the 50S stage provides an optimal balance for assessing sex variability and gametogenesis progression.

After selecting this stage, we performed a longitudinal experiment by separating sibling worms (DOB October 28, 2022) into individual cultures and assigning a unique ID to each worm. Two experimental groups were established: Control (uncut) and Regenerating (worms subjected to monthly amputation). Because worms ranged from 50 to 59 segments, amputation was performed approximately at mid-body. All worms were siblings derived from a single female and a single male mating. The amputation site was standardized between segments 25 and 26.

At the time of amputation, worms were relaxed in a 1:1 solution of 7.5% MgCl₂ in 0.2 μm–filtered artificial seawater (ASW). Following relaxation, worms were amputated and transferred to clean containers with ASW supplemented with penicillin–streptomycin (1:1000). Control worms were similarly transferred to clean containers with antibiotic-supplemented ASW. This procedure was repeated monthly at intervals of 30–45 days. Additionally, following each amputation event, the subsequent feeding was skipped for both control and regenerating worms, resulting in a starvation period of 5–7 days.

The anterior body was photographed at the start of the experiment (t0) and two months later (2 mpa), before amputation, to estimate age based on interocular distance (IOD) and the number of pigment spots on the second segment. For HCRs and gonial cluster scoring, amputated tails were collected at t0 and 2 mpa, and whole worms at 5 mpa.

The experiment began in 2023 with a total of 48 sibling 50S worms. Initially, worms were housed individually in 6-well plates (one worm per well), with each plate containing worms from a single condition. Because worms occasionally moved between wells, individuals found outside their assigned wells were removed from the experiment. To reduce experimental loss, worms that remained in their wells were transferred to 50 mL plastic containers after 1.5 months. In these plastic containers, a glass stone was placed at the bottom to facilitate tube construction in protected corners, reducing mortality associated with tube formation above the water line.

#### RNA-Sequencing of Gonial Clusters: Regeneration Experiment 4

We conducted an experiment similar to Experiment 1 using 50S worms derived from a single-pair mating. Worms were amputated once per month for two months, a duration previously shown to be sufficient to detect differences in interocular distance and pigment spot number. All worms were maintained under new moon conditions.

Two independent batches of worms were used: HE-03 (DOB February 6, 2024) and HE-05 (DOB March 10, 2025). The first was used for RNA-seq, and the second for validation of results using RT-qPCR. For each batch, 24-32 worms were included, with half assigned to the regenerating group and half to the control group. Surviving worms at the end of the experiment were collected and decerebrated for brain isolation.

Immediately after brain collection, worms were bisected, and the posterior body was dissected to collect coelomic fluid containing gonial clusters. Gonial cluster samples used for RNA isolation and sequencing were also used to determine sex via RT-qPCR, following a published protocol (dx.doi.org/10.17504/protocols.io.ewov1kq3kgr2/v1).

#### RNA-Sequencing of Heads: Regeneration Experiment 5

To characterize differential gene expression in heads across developmental stages, we performed RNA sequencing on heads collected from worms with 40–60 segments. The worms were collected during new moon conditions. We used several batches of worms with different dates of birth (DOB), transferring at least 10 worms from each batch into a 500 mL container before collecting them for the experiments. During collection, worms were amputated at the level of the peristomium, isolating the head and its appendages for RNA-seq. These heads were immediately fixed in TRIzol and RNA was isolated as described under RNA isolation. For the validation experiments, heads were collected and fast-frozen in liquid nitrogen and transferred to TRIzol during RNA isolation.

#### Regenerative response of brain: Regeneration Experiment 6

Sibling *Platynereis dumerilii* worms with 50–59 segments (DOB August 12, 2025) were used for this experiment. Animals were randomly assigned to two experimental groups (n = 5 worms per group): control condition (uncut) and 2 hpa (2-hour post-amputation). For the condition 2 hpa, worms were amputated transversely between segments 25 and 26, separating anterior (25 segments) and posterior body fragments (up to 30 segments). During the post-amputation period, all regenerating worms were maintained in artificial seawater (ASW) filtered through a 0.22 µm membrane and supplemented with penicillin–streptomycin (1:1000). Heads from animals in both conditions were collected simultaneously, 2 hours after amputation of the experimental group, and immediately flash-frozen in liquid nitrogen. The remaining anterior headless fragments and posterior body fragments from amputated worms, as well as the remaining body tissue from Control animals, were fixed overnight in 4% paraformaldehyde (PFA) prepared in phosphate-buffered saline (PBS) and filtered artificial seawater (ASW).

### Animal husbandry

In all experiments, worms were checked and fed twice per week, every Tuesday and Friday, with the standard *P. dumerilii* diet used in the Özpolat lab (a mixture of Spirulina and Sera Micron) (Kuehn et al., 2019). Seawater in single-worm cultures was changed every other week. The light regime corresponded to new moon conditions, with no nocturnal light exposure throughout the duration of the experiments. Each specimen used in experiments was labeled with the segment number, worm ID, condition, pigment score, interocular distance score, and date of birth whenever possible.

Single-worm cultures were fed twice weekly, receiving 1 mg of Spirulina and 0.3 mg of Sera Micron per worm at each feeding. This represents a slightly higher food intake than the standard feeding regimen described in Kuehn et al. (2019). To reduce allocation and handling bias, eligible worms were selected according to predefined experimental criteria and processed using balanced collection procedures. When worms were assigned to experimental groups, assignment was alternated between control and Highlander groups to maintain similar sample numbers across conditions. In addition, worm selection and cutting were separated between researchers whenever possible: one or two researchers isolated the worms, while another researcher performed the amputations without seeing the worms beforehand. This partial blinding reduced the possibility that knowledge of worm identity influenced the cutting procedure.

### Average lifespan account

A total of 71 batches of sibling worms were separated into LD boxes and monitored until all animals had either matured or died. Boxes with no remaining animals were marked as “gone.” For each fertilization batch, lifespan was defined as the day when the last animal in the box died and the box was empty. We measured the maximum lifespan for each box and calculated the mean, median, standard error, and standard deviation across the 71 batches. This assessment was performed with batches dated from July 27, 2021, corresponding to the earliest fertilization batch, until September 19, 2022. In total, 71 batches of sibling worms were followed from birth until death. Wild-type animals lived an average of 321.1 ± 132.2 days, with a median lifespan of 286 days and a range of 125–804 days.

### Pigment spot scoring

Pigment spot scoring was used to quantify pigmentation patterns in *P. dumerilii* as a marker of developmental age. We analyzed worms from experiments HE-01, HE-02, HE-04, and HE-05 at two time points: t0, before amputation, and 2 mpa. Each worm was assigned a unique ID, and information on segment number at t0, developmental condition, and date of birth was recorded. Images of the anterior body of each worm were captured at varying magnifications using a Zeiss Stemi 305 microscope, under consistent imaging conditions to minimize variation between samples. To ensure consistency, photos were taken at the same time across samples whenever possible.

All pigment scoring was completed using Fiji Image J software (Schindelin et al., 2012; Schneider et al., 2012). The worm images were imported into Fiji, and pigment spots were quantified specifically within the second segment with chaetae since this segment consistently displayed visible and reproducible pigmentation patterns across worms. The “Multi-Point” tool was then used to annotate each pigment spot. Each visible pigment spot within segment 2 was counted individually. Brown and red pigment spots, also known as chromatophores, were included when clearly distinguishable from the surrounding body.

After annotating each visible pigment spot, all selected points were added to the ROI Manager. The “Measure” function was then used to generate a spreadsheet containing the total number of annotated points. The final pigment count corresponded to the total number of pigment spots identified within the selected segment. To maintain consistency between samples, pigment spots were only counted when they appeared sufficiently opaque and visually distinguishable under the imaging conditions used.

Blurry or low-resolution images were excluded from the analyses. Annotated images and ROI files were further saved for documentation and reproducibility purposes. The complete protocol is available under (https://github.com/rannypribeiro/Ribeiro_et_al_Highlander.git).

### Interocular distance score (IOD)

The worms have two pairs of eyes, and we used the posterior pair to measure the interocular distance score (IOD). The same photos used to document pigment spots were used for interocular distance measurements, provided that they contained the second pair of eyes. We measured both control and highlander worms from experiments HE-01, HE-02, HE-04, and HE-05 at two time points: t0 and 2 mpa. In addition, worms from experiments HE-01 and HE-02 were measured at 5 mpa. Interocular distance (*a*) was measured as a straight line between the inner edges of the left and right eyes at the same horizontal level. Ocular distance (*b*) was measured from the leftmost edge of the left eye to the rightmost edge of the right eye at the widest horizontal point. Each measurement was recorded in a spreadsheet, and the ratios were calculated using a spreadsheet formula. The interocular distance score (IOD) was defined as the ratio of interocular distance to ocular distance (*a/b*)(Fig. S3E–O). We used this ratio because it provides a scale-independent metric, allowing comparison of images taken at different magnifications. Different magnifications were used to image the anterior body of the worms because image acquisition depended on worm size, with smaller worms requiring higher magnification. Due to these differences in magnification and inconsistencies in image resolution, absolute scale could not be reliably determined for all samples; therefore, ratios were used. All measurements were performed using Fiji/ ImageJ. The complete protocol is available at https://github.com/rannypribeiro/Ribeiro_et_al_Highlander.git.

### Whole-mount in situ hybridization chain reaction (HCR)

#### HCR probe design

We used our custom software (Kuehn et al., 2022) to synthesize HCR probes, targeting the following genes: *vasa*, estrogen receptor (*er-α*/ꞵ), methyl farnesoate receptor (*met*). The probes were designed to be compatible with selected HCR™ amplifiers (B1, B2, B3, B4) from Molecular Instruments, Inc (https://www.molecularinstruments.com/). Each probe set contained up to 32 pairs of 25 bp sequences. The designed probe sequences were sent to Integrated DNA Technologies (https://www.idtdna.com) for synthesis at 5 nmol per probe set (containing up to 32 pairs of sequences). Before use, probes were re-suspended to 50 µM with Nuclease Free water (Invitrogen UltraPureTM Distilled Water, 10977-015).

#### Tissue collection and HCR procedures

We performed hybridization chain reaction (HCR) RNA detection experiments in tails, trunks, or whole worms depending on the experimental design (see above). HCR was performed following previously described protocols (Kuehn et al., 2022) with the following modifications: with two modifications: (1) proteinase K treatment was replaced with a Tween-20-based permeabilizing solution, and (2) some samples were preserved in High SDS Buffer instead of methanol.

Samples were collected and anesthetized in a 1:1 solution of 0.22 µm filtered natural seawater (FNSW) and 7.5% MgCl₂ for 15–30 minutes, followed by overnight fixation in 4% paraformaldehyde (PFA). The fixative was prepared at a 1:2:1 ratio of 16% PFA, 2× phosphate-buffered saline (PBS), and filtered artificial seawater (ASW).

Following fixation, samples were processed using one of two preservation methods, depending on the experiment: (1) dehydration through a graded methanol series (25%, 50%, 75%, and 100% methanol; 10–30 minutes per step), or (2) direct incubation in High SDS Hybridization buffer. After these procedures, samples were stored at -20 °C in either 100% methanol or High SDS Hybridization buffer until further use. Methanol dilutions were prepared in 1× PBS containing 0.1% Tween-20 (1× PBSt). All PBS solutions were prepared with diethyl pyrocarbonate (DEPC)-treated, ultrafiltered water.

Samples preserved in High SDS Hybridization buffer were washed after fixation as follows: two washes in PBS for 5 minutes each, followed by one wash for 30 minutes. Samples were then washed three times for 20–30 minutes in a Tween-20-based detergent solution, incubated in High SDS Hybridization buffer for 30 minutes at 37 °C, and stored at −20 °C until continuation of the HCR protocol. Detection steps were performed as described for methanol-preserved samples (Kuehn et al., 2022).

The probe hybridization buffers, wash buffers, amplification buffers and HCR™ amplifiers were obtained from Molecular Instruments, Inc (https://www.molecularinstruments.com/). Samples were mounted on cover slides with SlowFade™ Glass Soft-set Antifade Mountant, with DAPI (Invitrogen™ S36920) or SouthernBiotech™ Fluoromount-G™ Slide Mounting Medium (SouthernBiotech™ 010001).

### RNA sequencing experiments

#### RNA isolation for RNA-sequencing

Two independent RNA-seq experiments were performed using different tissues and RNA-isolation procedures. For the head RNA-seq experiment, freshly collected heads were immediately transferred to TRIzol reagent and stored at −80 °C until processing. Total RNA was extracted from the TRIzol-preserved samples and further purified using the RNeasy Mini Kit (Qiagen). For the gonial-cluster RNA-seq experiment, coelomic contents enriched in gonial clusters were collected as described previously (Ribeiro et al., 2025). RNA was isolated using the Arcturus PicoPure RNA Isolation Kit (Applied Biosystems). For both experiments, RNA concentration and purity were assessed using a NanoDrop One-C spectrophotometer. Samples submitted for sequencing had RNA concentrations of approximately 10 ng/µL and A260/A280 ratios between 1.8 and 2.0.

#### RNA sequencing

At the sequencing facility (GTAC, WashU), total RNA integrity was determined using an Agilent Bioanalyzer. Samples with clean, intact ribosomal peaks and at least 15ng of total RNA were selected for sequencing. ds-cDNA was prepared using the SMARTseq mRNA kit (Takara-Clontech #634773) per manufacturer’s protocol. cDNA was fragmented using a Covaris E220 sonicator using peak incident power 18, duty factor 20%, cycles per burst 50 for 120 seconds. cDNA fragments were blunt-ended, adenylated at their 3′ ends, and ligated to Illumina sequencing adapters. Ligated fragments were then amplified for 15 cycles using primers incorporating unique dual index tags. Fragments were sequenced on an Illumina NovaSeq X Plus using 150-base paired-end reads.

#### Bioinformatic analyses

Raw sequencing reads were initially assessed using FastQC(Andrews, 2010). Adapter sequences and low-quality bases were removed using Trimmomatic (Bolger et al., 2014), after which read quality was reassessed with FastQC. Only libraries showing satisfactory overall sequence quality and no detectable adapter contamination were retained for downstream analysis.

The gonial-cluster and head RNA-seq datasets were processed separately using experiment-specific reference transcriptomes. Reads from coelomic samples enriched in gonial clusters were analyzed against the reference transcriptome described by Ribeiro et al. (2025) using Trinity RNA-seq (Grabherr et al., 2013; Haas et al., 2013). Reads from head samples, which were used to identify age-associated transcriptional changes, were analyzed against the reference transcriptome derived from the genome assembly published by Mutemi et al. (2025).

As an additional quality-control step, reads from each library were mapped against the corresponding reference transcriptome, and mapping statistics were compared across biological replicates. Libraries with substantially different mapping performance were excluded.

Following quality-control filtering, only the 40-segment and 60-segment head samples were included in the differential expression analysis of the head RNA-seq experiment. Only two 50-segment samples passed quality control, which was below the predefined minimum of three biological replicates per group. Therefore, 40-segment and 60-segment worms were compared using six biological replicates per group.

For the gonial-cluster RNA-seq experiment, reads from both Control and Highlander samples mapped well to the reference transcriptome. However, differential expression analysis was restricted to females, for which five Control and four Highlander biological replicates were available. Males could not be compared because only two Control males were identified, which was below the minimum number of replicates required for analysis. Because the sex of the worms was unknown at the time of sample collection, all 15 samples from each experimental condition were submitted for sequencing. An aliquot of RNA from each sample was retained for sex identification by RT-qPCR using our previously established protocol available on protocols.io (dx.doi.org/10.17504/protocols.io.ewov1kq3kgr2/v1). Subsequent sexing revealed that only two of the Control samples were male.

For both experiments, differential expression analyses were performed using the Trinity v2.8 differential expression workflow. Transcript abundance was estimated for each sample using Kallisto (Bray et al., 2016), and differential expression was assessed in R using DESeq2 (Love et al., 2014). Expression profiles were compared at the ‘Trinity gene’ level. Differential expression results and functional enrichment analyses are available in our GitHub repository (https://github.com/rannypribeiro/Ribeiro_et_al_Highlander). The repository also includes two Jupyter notebooks and accompanying scripts designed to facilitate data exploration, generate heatmaps, and visualize differential gene expression patterns.

### RT-qPCR

#### RNA isolation for RT-qPCR analysis

Total RNA was extracted from samples using protocols adapted to the tissue type. For head samples subjected to RT-qPCR analysis, RNA was extracted using the column-based RNeasy Mini Kit (Qiagen). RNA quality and concentration were assessed prior to reverse transcription.

Complementary DNA (cDNA) was synthesized from RNA templates using the following kits according to the manufacturers’ instructions: First Strand cDNA Synthesis Kit for RT-PCR (11483188001, Roche), High-Capacity cDNA Reverse Transcription Kit with RNase inhibitor (4374966, ThermoFisher Scientific), High-Capacity RNA-to-cDNA™ Kit (4387406, ThermoFisher Scientific).

#### RT-qPCR setup and cycling conditions

Quantitative PCR reactions were performed in 20 µL volumes using a SYBR Green detection system. Each reaction contained 2–4 µL of cDNA template, 1.25 µL of forward primer, 1.25 µL of reverse primer, SYBR Green master mix (SsoAdvanced SYBR Green, Bio-Rad), and nuclease-free water to a final volume of 20 µL. Each sample was run in at least two technical replicates per gene.

Reactions were loaded in duplicate into 96-well plates and run on a Bio-Rad CFX Opus 96 using the following cycling conditions: 95 °C for 2 min (initial denaturation); 39 cycles of 95 °C for 10 s (denaturation) and 60 °C for 30 s (annealing/ extension), and a final step for melt curve analysis, 65 °C to 95 °C with 5-second increments.

Fluorescence data were collected using SYBR Green chemistry. Quantification cycle (Cq) values were obtained using the instrument software. Melt curve analysis was performed for each reaction to verify amplification specificity. Reactions showing multiple melting peaks or abnormal amplification profiles were excluded. Negative controls (no-template controls) were required to show no amplification or Cq values >32–39.

#### Reference gene validation and normalization

To identify suitable reference genes, the expression stability of candidate housekeeping genes was evaluated across all samples. A suitable reference gene was defined as one showing minimal variation in Cq values and little or no systematic association with biological or technical variables. Linear models were therefore used to assess whether Cq values varied with segment number, sex, or experimental batch. Stability was further evaluated using variance-based measures, including the standard deviation of Cq values and model residual error. Genes with the lowest overall variability and the weakest dependence on these variables were considered the most stable. Based on these analyses, cdc5 was selected as the reference gene for RT-qPCR analyses of head samples, whereas 18S and axin were selected as reference genes for RT-qPCR analyses of RNA isolated from coelomic contents.

#### RT-qPCR data analysis

All analyses were performed in R. Technical replicates were averaged prior to downstream analysis. Samples with inconsistent replicates or poor amplification quality were excluded. Statistical analyses included linear modeling to assess reference gene stability and visualization of Cq distributions across experimental variables.

### Morphological staging of female gonial clusters

#### Gonial cluster scoring

Gonial clusters were scored in whole-mount hybridization chain reaction (HCR) specimens. Worms were fixed and processed as described in the section “Whole-mount HCR.” Following HCR, specimens were divided into anatomically ordered body fragments to permit flat mounting and imaging of the entire trunk. Fragments were mounted in SlowFade™ Gold Antifade Mountant with DAPI on coverslips and imaged using a Zeiss LSM 900 confocal microscope equipped with a 40× objective. Overlapping tiled z-stacks were acquired to cover the width and depth of each body fragment where specimen geometry permitted. Z-stacks were acquired at 8 µm intervals along the depth, depending on sample thickness, covering about 60–250 µm inside the worm. Stage assignments were made by examining the complete z-stack rather than individual optical planes or maximum-intensity projections.

After imaging, files were assigned randomized names using a custom Python script so that sample identities were concealed during scoring. Imaging parameters, fragment position, number of body segments, and gonial-cluster stages were recorded for each fragment.

Germline structures were identified using HCR detection of *vasa* transcripts in combination with DAPI-stained nuclear morphology and anatomical position. Thus, *vasa* served as an internal germline marker rather than an experimental positive control. Discrete *vasa*-positive cell groups within the trunk coelom were classified as gonial clusters. The anterior germline cluster was excluded; only gonial clusters distributed throughout the trunk were included in the analysis.

Female gonial clusters were staged according to the morphological framework for early oogenesis in *Platynereis dumerilii* described by Fischer (1974, 1975), using the reference examples shown in Fig. 2B and Supplementary Fig. S3A. We grouped Stages 1 and 2 into a single Stage 1/2 category because these early stages are difficult to distinguish reliably in whole-mount HCR samples. Stage 3 was characterized by expansion of the nucleolus. Stage 4 was characterized by the appearance of yolk droplets and enlargement of the nucleus and was further divided into Stage 4-cluster, representing earlier Stage 4 oocytes that remained organized within clusters, and Stage 4-singlet, representing later Stage 4 oocytes that were no longer associated with a cluster (Fig. 2B).

Because early oogenesis can be asynchronous within individual clusters, a single cluster frequently contains cells representing more than one morphological stage. Each cluster was therefore assigned a maximum qualifying stage, defined as the most advanced stage represented by at least three unambiguously identifiable cells. For example, a cluster containing at least three Stage 4 cells was classified as Stage 4-cluster, even if cells at Stages 2 and 3 were also present. In contrast, if only one or two Stage 4 cells were present, the cluster was assigned to the next most advanced stage represented by at least three cells. Isolated Stage 4 oocytes were recorded separately as Stage 4-singlets.

Clusters with insufficient signal, incomplete z-coverage, ambiguous cell boundaries, or truncation at a specimen-cutting boundary were classified as unscorable and excluded from stage-frequency analyses. Representative annotated examples and the complete scoring dataset, including a custom Python script used to randomize images, are provided in our GitHub repository (https://github.com/rannypribeiro/Ribeiro_et_al_Highlander).

### Statistical analyses

#### Lifespan and maturation curves

Analyses were performed separately for experiments, comparing Control and Highlander animals. Lifespan and maturation ages were calculated as days from birth to the recorded death or maturation date; animals were not filtered by developmental stage or initial segment count. Animals with missing dates were excluded from the corresponding analysis rather than treated as censored, and animals with unknown sex were excluded only from sex-based analyses. Survival and time-to-maturation curves were estimated with the Kaplan–Meier method, and conditions were compared using log-rank tests; plotted medians are medians of observed event ages. Within each experiment and condition, the association between initial segment number and maturation age was tested by ordinary least-squares linear regression. Lifespan and maturation ages were also analyzed using Gaussian linear models with condition, sex, and their interaction, followed by Tukey-adjusted pairwise comparisons of estimated marginal means. Daily juvenile, dead-before-maturity, and mature percentages were descriptive and were calculated for animals with known death dates.

#### Gonial cluster staging

Gonial-cluster stage distributions were analyzed using negative-binomial generalized linear mixed-effects models, with gonial-cluster count as the response variable, condition (Highlander and Control) or time (t0, 2 mpa), stage, and their interaction as fixed effects. Post hoc contrasts of estimated marginal means were used to compare conditions or time points within each developmental stage. Pearson chi-square tests were additionally used to assess overall differences in aggregated stage distributions. Changes in the total number of gonial clusters between t0 and 2 mpa in Highlander worms were also assessed using a two-sided Wilcoxon rank-sum test because the individuals sampled at the two time points were independent. Data were summarized and displayed as individual worms with the median and standard deviation.

#### Pigment spot analyses

Pigment spot number was analyzed longitudinally to test whether pigment accumulation over time differed between conditions. For visualization, Pigments Added was calculated for each animal as the paired difference between pigment number at 2 months post-amputation (2 mpa) and baseline (t0). Analyses were performed for the pooled cohort, separately for 50S and 60S+ animals, and separately by experiment.

Pigment measurements were modeled using a Gamma generalized linear mixed-effects model with a log link. The model included fixed effects for condition, time point, and their interaction, with a random intercept for worm ID: value ∼ condition * measurement + (1 | worm_id). The primary inferential test was the condition-by-timepoint interaction, which tests whether the longitudinal change in pigment number differed between conditions. These interaction p-values are the p-values shown in the figures; testing differential change over time.

#### IOD score analyses

Interocular distance (IOD) was measured as the distance between the second pair of eyes in *P. dumerilii*. Because absolute metric scaling was not available for all images, IOD was normalized within each image as the ratio (*a/b*), where *a* represents the distance between the inner edges of the posterior pair of eyes and *b* represents the distance between their outermost edges, measured along the same horizontal axis. IOD was analyzed longitudinally using animals with available baseline (t0) and follow-up (2 mpa or 5 mpa) measurements. Fold change was calculated as follow-up IOD divided by baseline IOD, and the natural logarithm of fold change was used for statistical inference. Within each condition, two-sided one-sample t-tests tested whether the mean log fold change differed from zero. This is equivalent to a paired t-test comparing log-transformed follow-up IOD with log-transformed baseline IOD. Geometric mean fold change was calculated as exp(mean log fold change), and 95% confidence limits were calculated by exponentiating the t-based confidence limits from the log fold-change scale. Tests and summaries were calculated overall for both 2 mpa and 5 mpa, and separately by experiment for 2 mpa. These analyses evaluated within-condition longitudinal changes only.

#### Analyses of number of segments in amputated tails

The number of segments in amputated tails was analyzed longitudinally using linear mixed-effects models fitted by maximum likelihood. Worm identity was included as a random intercept to account for repeated measurements of the same individual. Equivalent models were fitted for tail size expressed as a percentage of the worm’s initial segment number. Estimated marginal means were used for post hoc comparisons among time points and for comparisons of each time point with baseline (0 months post-amputation); *P* values were adjusted for multiple testing using the Benjamini–Hochberg procedure. The association between initial worm size and regenerated-tail size was examined using a mixed-effects model adjusted for time and, separately, by Spearman’s rank correlation using each worm’s latest post-amputation measurement. As a nonparametric sensitivity analysis, changes across time were evaluated with a Friedman test restricted to worms with complete observations, i.e., worms that survived until 7 mpa. Initial segment numbers were compared among experiments using a Kruskal–Wallis test. Individual longitudinal trajectories were additionally summarized using worm-specific linear regressions. All tests were two-sided.

### Figure editing

Figures were edited using Adobe Illustrator version 28.6. Clip art was obtained from open-access resources, including NIH BioArt (https://bioart.niaid.nih.gov/) and Open Clipart (https://openclipart.org/). Confocal and light microscopy images were edited in Fiji/ Image J. Life stage cartoons were created by RPR using Procreate version 5.4.12.

## Supplementary Figures

**Figure S1.**
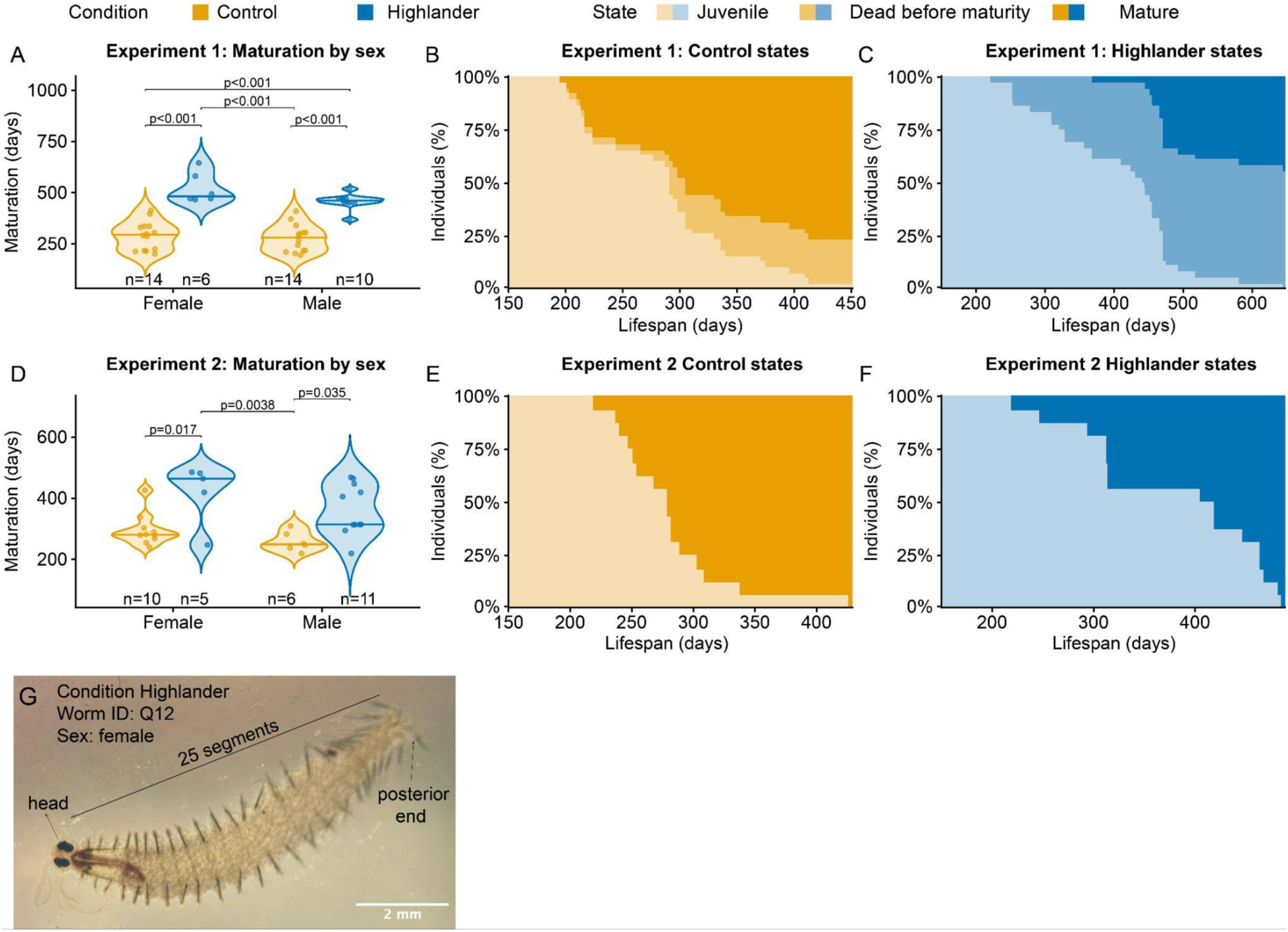
Maturation by sex and lifespan states by experiment. (A–C) HE-02: maturation by sex (A) and individual states over time for Control (B) and Highlander animals (C). (D–F) HE-04: maturation by sex (D) and individual states over time for Control (E) and Highlander animals (F). Individual states were classified as juvenile, dead before maturity, or mature and are shown using condition-specific orange and blue fills. These plots account for mortality occurring before sexual maturation. In HE-02, mortality before maturity was 23.6% in Controls and 55% in Highlander animals, whereas no mortality before maturity occurred in HE-04. (G) Light microscopy images of a female Highlander specimen. Note that the posterior end of this individual failed to regenerate the pygidium (tail).

**Figure S2.**
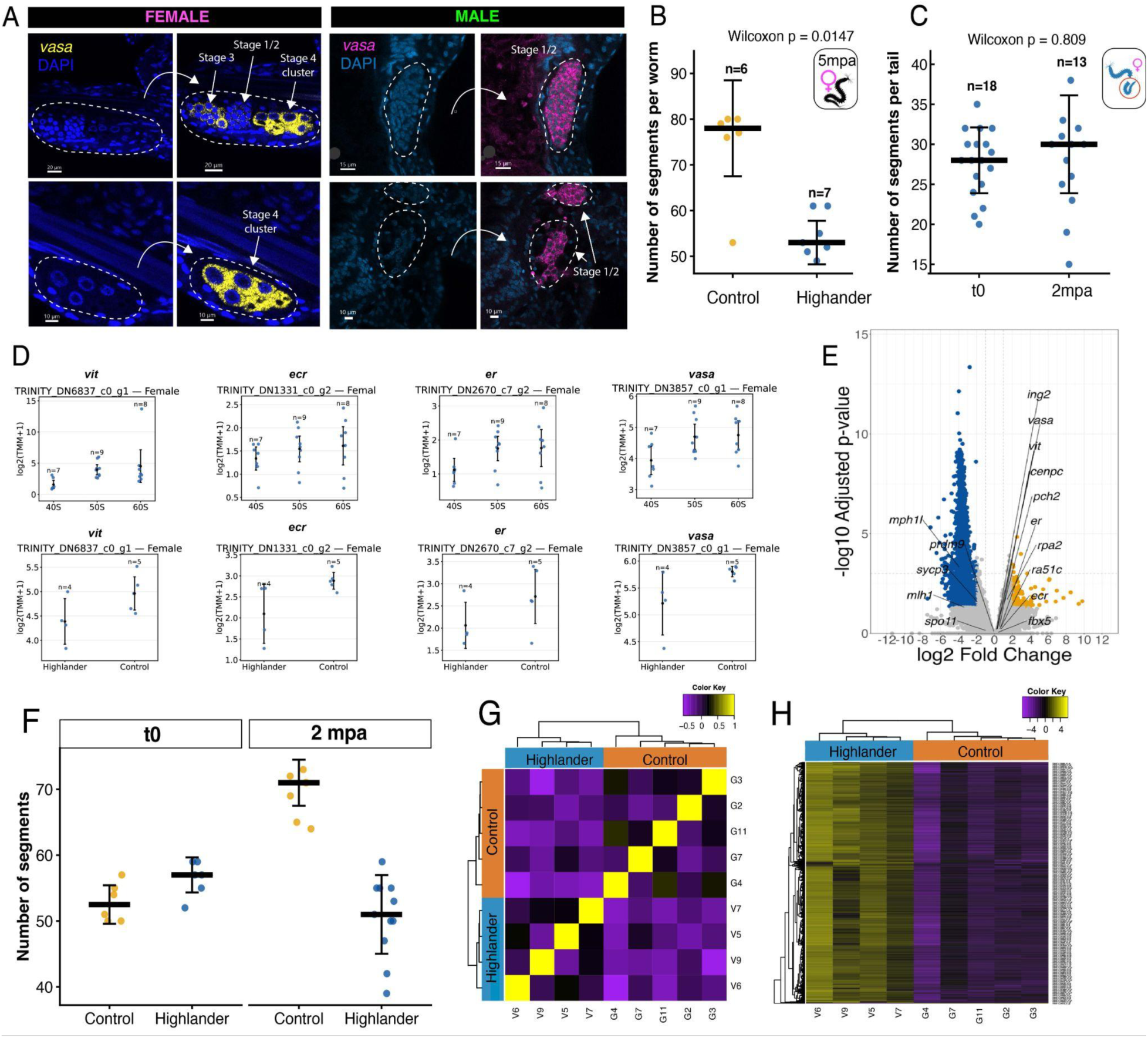
Regeneration delays gametogenesis and sexual maturation. (A–C) Results from Experiment 3. (A) Examples of gonial cluster stages in females and males. (B) Differences in worm size at 5 months post-amputation (mpa). (C) Size comparison between the original amputated tails at t0 and regenerated tails at 2 mpa in Highlander worms. At t0, an average of 28 segments were removed, whereas at 2 mpa, worms had regenerated an average of 30 segments following the second amputation. (D–H) Results from Experiment 4. (D) Comparison of RNA-seq gene-expression profiles from the reference dataset of unamputated wild-type (WT) *P. dumerilii* with 40, 50, and 60 segments (Ribeiro et al., 2025) and Control and Highlander worms from this study; selected genes: *vit*, *ecr*, *er*, and *vasa*. (E) Volcano plot highlighting selected gametogenesis-related genes (*p* < 0.05). (F) Number of segments in Control and Highlander worms at t0 and 2 mpa. (G) Quality-control analysis supporting the differential-expression (DE) analysis: sample-to-sample Pearson correlation matrix showing clustering of biological replicates. (H) Heatmap of differentially expressed genes between Highlander and Control females.

**Figure S3.**
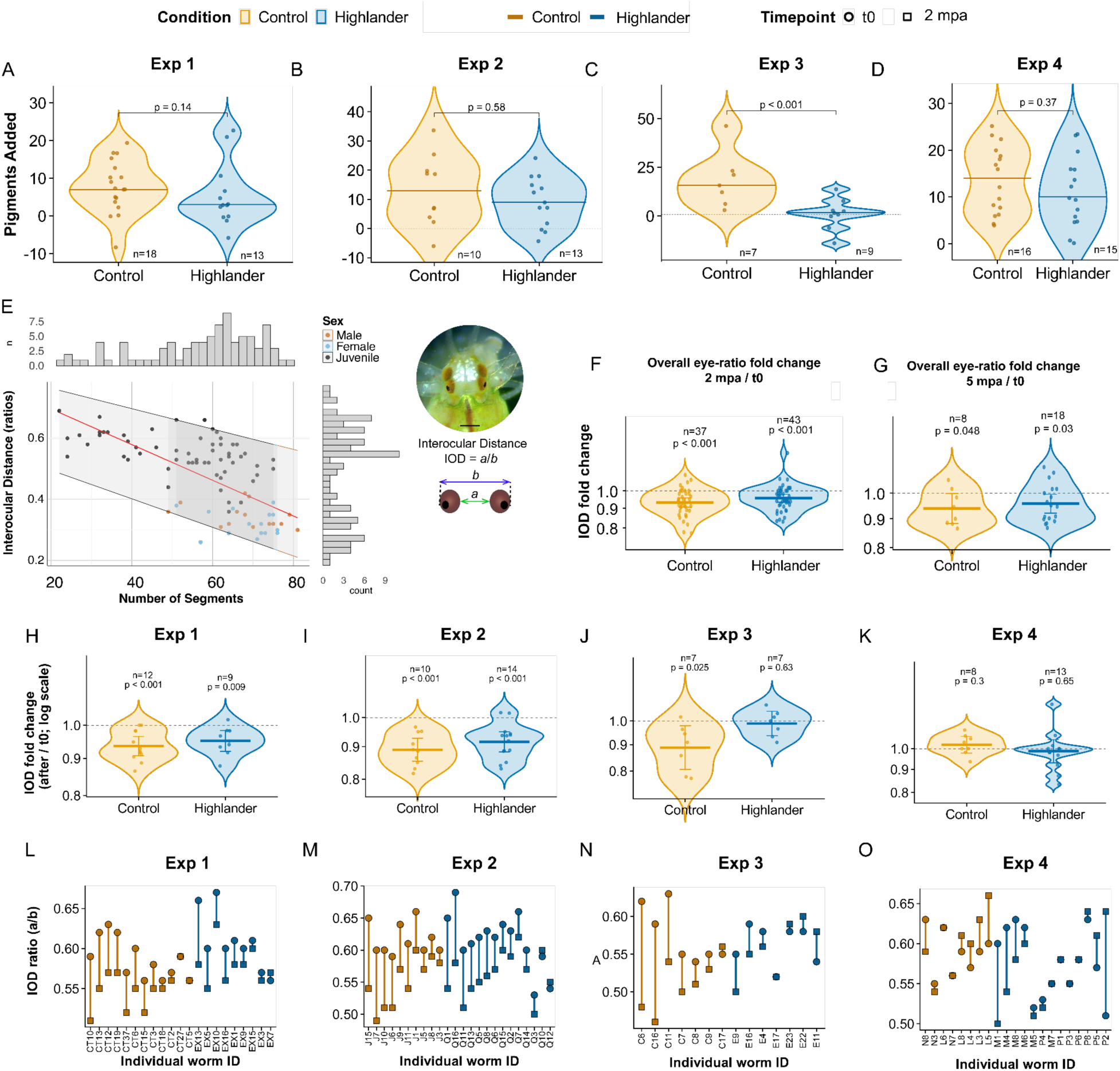
Pigment addition and interocular distance (IOD) across experiments. (A–D) Pigment addition across Experiments 1–4; violin plots show individual animals, group medians, and a zero-change reference line. (E) IOD across developmental stages and a light microscopy image of the head of a juvenile *P. dumerilii* (left); scale bar, 200 μm. (F–K) IOD fold changes for Control and Highlander worms. Violin plots show individual animals, geometric means, and a no-change reference line at 1, with fold change displayed on a logarithmic axis. (F) Overall change at 2 mpa and (G) at 5 mpa. (H–K) IOD fold changes across Experiments 1–4. (L–O) Changes in IOD for individual worms in Experiments 1–4. Plots show individual animals, geometric means, and 95% confidence intervals, with a logarithmic fold-change axis and a no-change reference line at 1. IOD is used as a proxy for eye size, based on the assumption that eye growth is accompanied by a decrease in the distance between the eyes.

**Fig. S4.**
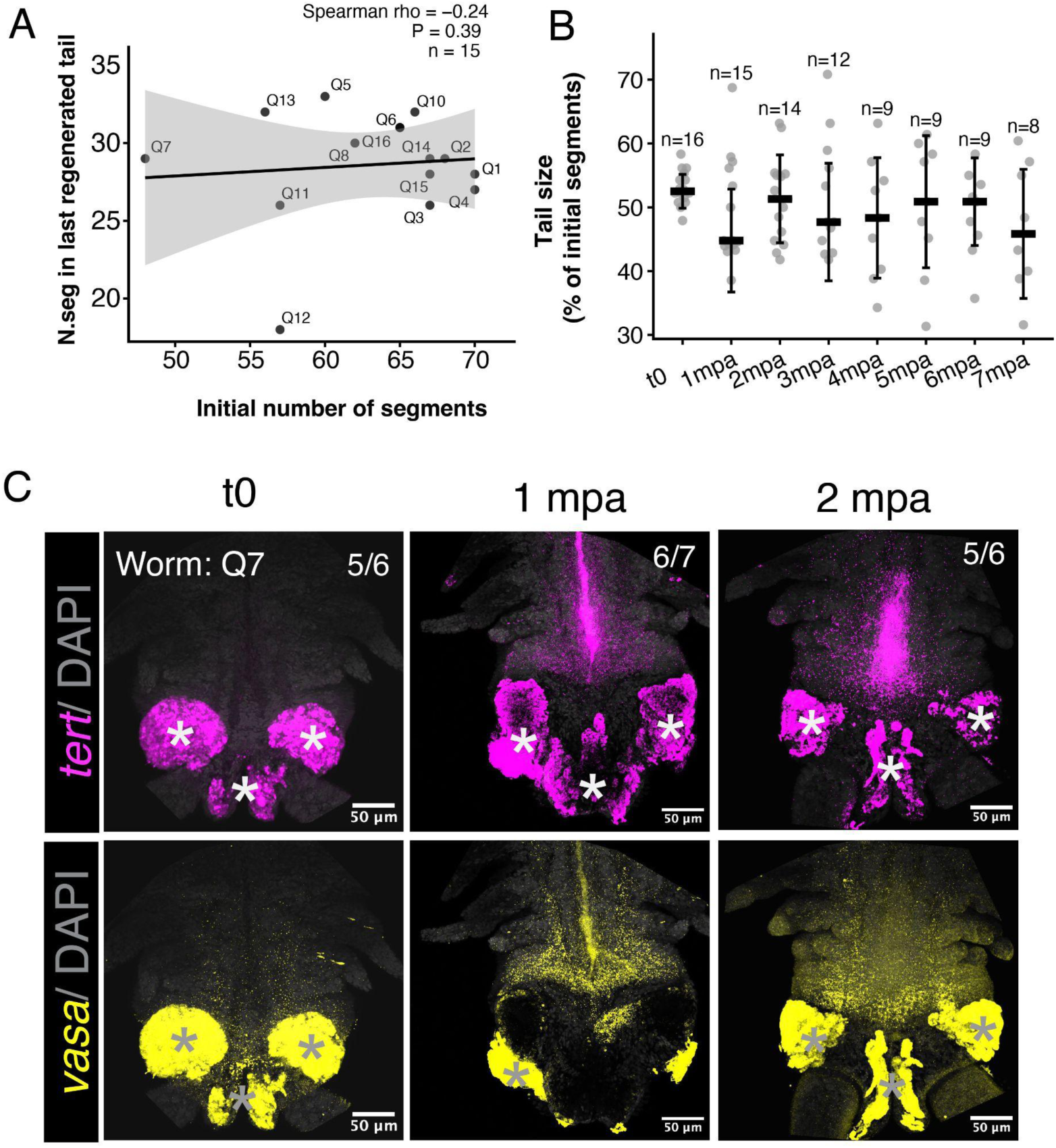
Measurements of amputated tails in Experiment 2 and corresponding HCR images. (A) Number of segments in the last regenerated tail collected from Highlander worms. Because not all worms survived through 7 mpa, the last tail corresponds to different amputation cycles among individuals. (B) Percentage of total body size represented by the amputated tail. Because the experimental design involved amputation at approximately half the body length, values near 50% indicate that the amputated tail represented approximately half of the worm in terms of segment number. (C) Full images of tails collected from worm Q7, highlighted in Fig. 4. Glands in the pygidium, the posteriormost region of the tail, show autofluorescence (asterisks); Fig. 4 therefore shows only the posterior growth zone (PGZ).

**Figure S5.**
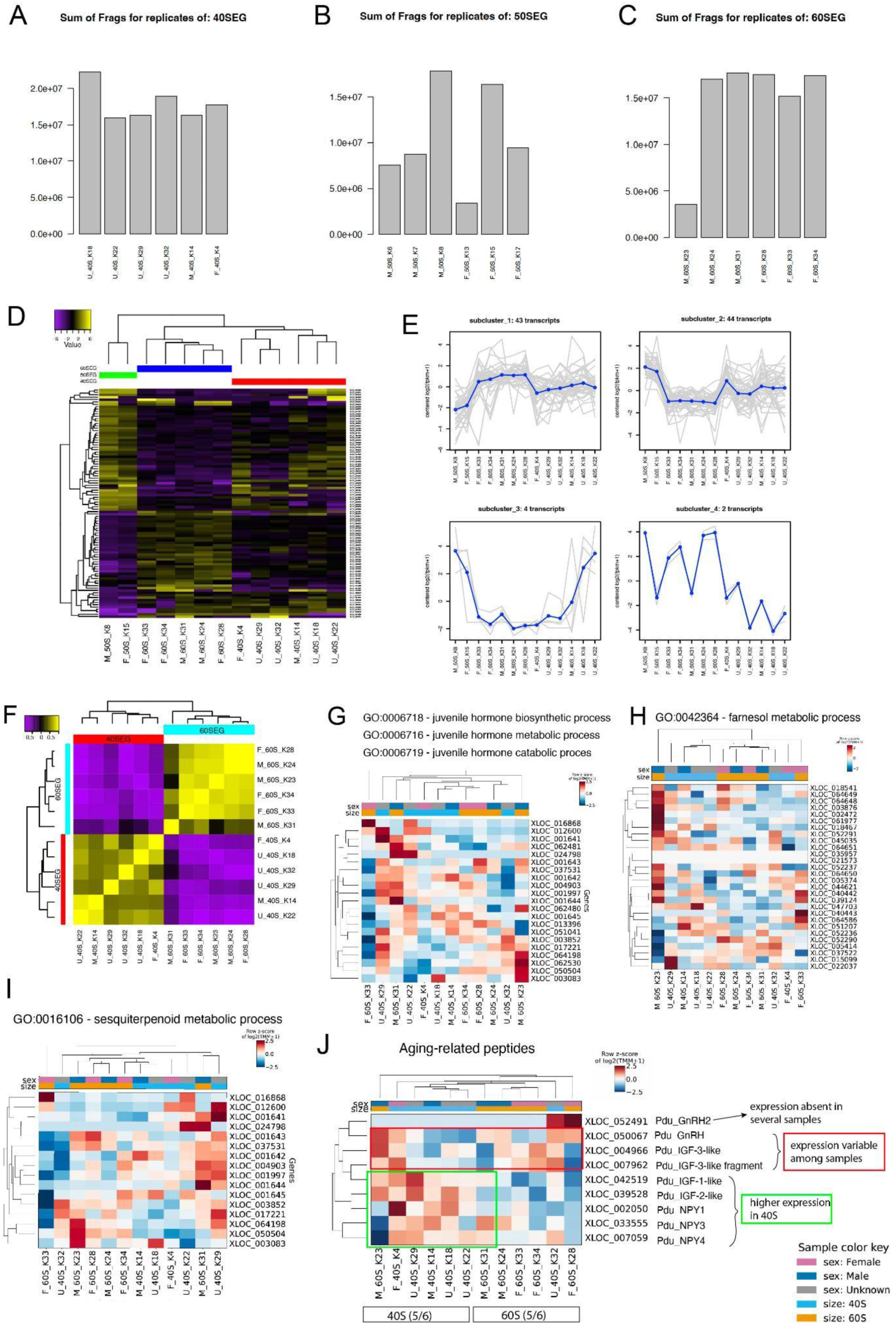
Quality control and exploratory analysis of head RNA-seq samples. (A– C) Summary of sequencing reads mapped to the reference *Platynereis dumerilii* transcriptome across developmental-age groups. (D–E) Unbiased differential-expression analysis including all biological replicates that passed initial quality-control criteria, showing differences in mapping and transcript abundance among samples. Samples from the 50-segment (50S) group displayed greater variability in their mapping profiles than the 40S and 60S samples and were therefore excluded from the final differential-expression comparison. (F) Sample-to-sample Pearson correlation matrix generated using the Trinity differential-expression workflow for the retained 40S and 60S samples. The correlation matrix was used as an additional quality-control assessment to evaluate the similarity and clustering of biological replicates within each developmental group. (G–J) Expression of genes associated with Gene Ontology (GO) terms related to juvenile hormone and methyl farnesoate pathways in the head RNA-seq dataset.

